# Reliable protein-protein docking with AlphaFold, Rosetta, and replica-exchange

**DOI:** 10.1101/2023.07.28.551063

**Authors:** Ameya Harmalkar, Sergey Lyskov, Jeffrey J. Gray

## Abstract

Despite the recent breakthrough of AlphaFold (AF) in the field of protein sequence-to-structure prediction, modeling protein interfaces and predicting protein complex structures remains challenging, especially when there is a significant conformational change in one or both binding partners. Prior studies have demonstrated that AF-multimer (AFm) can predict accurate protein complexes in only up to 43% of cases.^1^ In this work, we combine AlphaFold as a structural template generator with a physics-based replica exchange docking algorithm to better sample conformational changes. Using a curated collection of 254 available protein targets with both unbound and bound structures, we first demonstrate that AlphaFold confidence measures (pLDDT) can be repurposed for estimating protein flexibility and docking accuracy for multimers. We incorporate these metrics within our ReplicaDock 2.0 protocol^2^ to complete a robust in-silico pipeline for accurate protein complex structure prediction. AlphaRED (AlphaFold-initiated Replica Exchange Docking) successfully docks failed AF predictions including 97 failure cases in Docking Benchmark Set 5.5. AlphaRED generates CAPRI acceptable-quality or better predictions for 63% of benchmark targets. Further, on a subset of antigen-antibody targets, which is challenging for AFm (20% success rate), AlphaRED demonstrates a success rate of 43%. This new strategy demonstrates the success possible by integrating deep-learning based architectures trained on evolutionary information with physics-based enhanced sampling. The pipeline is available at github.com/Graylab/AlphaRED.

## Introduction

In-silico protein structure prediction, *i*.*e*., sequence to structure, tackles one of the core questions in structural biology. AlphaFold^3^ has brought a paradigm shift in the field of structural biology by intertwining deep-learning (DL) tools with evolutionary data to predict single-chain structures with high accuracy. Further, AlphaFold-multimer^4^ (AFm) and related work^5,6^ have demonstrated the utility of AlphaFold to predict protein complexes. The association of proteins to form transient or stable protein complexes often involves binding-induced conformational changes. Capturing conformational dynamics of protein-protein interactions is another grand challenge in structural biology, and many physics-based (computational) approaches have been used to tackle this challenge.^2^ Computational tools have sampled the uncharted landscape of protein-protein interactions by emulating kinetic mechanisms such as conformer selection and induced-fit and by identifying energetically stable binding states. However, these tools are hampered by the accuracy of the energy functions and the limitations of time and length scales for sampling. In fact, AF-multimer (AFm) predicted accurate protein complexes in only 43% of cases in one recent study.^1^ As the development of DL-based tools have unveiled ground-breaking performance in structure prediction, integration of a biophysical context has potential to strengthen prediction of protein assemblies and binding pathways.

Blind docking challenges prior to AF, particularly CASP13-CAPRI and CASP14-CAPRI experiments, reported high-quality predictions for only 8% targets.^7,8^ With the availability of AF and AFm, the CASP15-CAPRI experiment stood as its first blind assessment for prediction of protein complexes and higher-order assemblies.^9^ In this round, the docking community relied on AF and AFm for single-structure or complex predictions. Given that AlphaFold generates a static three-dimensional structure, it has been unclear whether conformational diversity could be captured by AlphaFold. In other terms, given a protein sequence, could AlphaFold generate ensembles of structures that include both unbound and bound conformations? Additionally, can AlphaFold reveal intrinsic conformational heterogeneity?

To diversify model complexes generated with AlphaFold-multimer in the recent round of CASP15, predictors employed tuning parameters such as dropout,^10^ higher recycles on inference^11^ or modulating the MSA inputs^12,13^ with the amino acid sequence. While these approaches demonstrated the ability to generate broader conformational ensembles, AFm performance still worsens with a higher degree of conformational flexibility between unbound and bound targets.^1^ Prediction accuracies especially deteriorated in bound complex regions involving loop motions, concerted motions between domains, rearrangement of secondary structures, or hinge-like domain motions, *i*.*e*., large-scale conformational changes, which are also challenging for conventional docking methods.^14^

Unlike state-of-the-art docking algorithms, AlphaFold’s output models incorporate a residue-specific estimate of prediction accuracy. This suggests a few interesting questions: (1) Do the residue-specific estimates from AF/AFm relate to potential metrics demonstrating conformational flexibility? (2) Can AF/AFm metrics deduce information about docking accuracy? (3) Can we create a docking pipeline for in-silico complex structure prediction incorporating AFm to convert sequence to structure to docked complexes?

Recent work in physics-based docking approaches tested induced-fit docking^2^, large ensembles^15^, and fast-fourier transforms with improved energy functions^16^ to capture conformational changes and better dock protein structures. Coupling temperature replica exchange with induced-fit docking, ReplicaDock 2.0^2^ achieved successful local docking predictions on 80% of rigid (unbound-to-bound root mean square deviation, RMSD_UB_ < 1.1Å) and 61% medium (1.1 ≤ RMSD_UB_ < 2.2 Å) targets in the Docking Benchmark 5.0 set^17^. However, like most state-of-the-art physics-based docking methods, ReplicaDock 2.0 performance was limited for highly flexible targets: 33% success rate on targets with RMSD_UB_ ≥ 2.2 Å. Promisingly, by focusing backbone moves on known mobile residues (*i*.*e*., residues that exhibit conformational changes upon binding), ReplicaDock 2.0 sampling substantially improved the docking accuracy. But the flexible residues must first, somehow, be identified. Additionally, physics-based docking is quite slow (6-8 hrs on a 24-core CPU cluster) compared to recent DL based docking tools (0.1-10 minutes on a single NVIDIA GPU). However, docking-specific DL tools such as EquiDock^18^ and dMASIF^19^ do not allow for protein flexibility, and recent tools like GeoDock,^20^ and DockGPT^21^ have very limited backbone flexibility. Further, all of these DL docking tools have low success rates on unbound docking targets such as those in Docking Benchmark 5.5.^20^

In this work, we combine the features of a top deep learning approach (AlphaFold-multimer^4^) with physics-based docking schemes (ReplicaDock 2.0^2^) to systematically dock protein interfaces. The overarching goal is to create a one-stop, fully automated pipeline for simple, reproducible, and accurate modeling of protein complexes. We investigate the aforementioned questions and create a protocol to resolve AFm failures and capture binding-induced conformational changes. We first assess the utility of AFm confidence metrics to detect conformational flexibility and binding site confidence. Next, we feed these metrics and the AFm-generated structural template to ReplicaDock 2.0, creating a pipeline we call AlphaRED (AlphaFold-initiated Replica Exchange Docking). We test AlphaRED’s docking accuracy on a curated set of benchmark targets of bound and unbound protein structures of varying levels of binding-induced conformational change, including antibody-antigen interfaces, which additionally challenge AF2m due to the lack of evolutionary information across the interface.^22,23^ In summary, we to assess the promise of combining the best of deep learning and biophysical approaches for predicting challenging protein complexes.

## Results

### Dataset curation

We curated a dataset for conformational flexibility from the Docking Benchmark Set 5.5 (DB5.5)^17^, which comprises experimentally characterized (X-ray or cryo-EM) structures of bound protein complexes and their corresponding unbound protein subunits. Each protein target (with unbound and bound structures) is classified based on their unbound-to-bound root-mean-square-deviation (RMSD_UB_) as rigid (RMSD_UB_ ≤ Å), medium (1.2 Å < RMSD_UB_ ≤ 2.2 Å) or difficult (RMSD_UB_ ≥ 2.2 Å). Furthermore, due to the poor performance of AlphaFold and other predictor groups in predicting antibody-antigen targets in the recent CASP15-CAPRI round^24^, we identified a subset comprising only antibody-antigen complexes (including single-domain antibodies, or nanobodies) extracting all 67 antibody-antigen structures from the DB5.5^17,25^ set. The comprehensive dataset includes 254 protein targets exhibiting binding-induced conformational changes.

For each protein target, we extracted the amino acid sequences from the bound structure and predicted a corresponding three-dimensional complex structure with the ColabFold implementation (github.com/YoshitakaMo/localcolabfold) of the AlphaFold multimer v2.3.0 (released in March 2023) for the 254 benchmark targets from DB5.5. Being trained on experimentally characterized structures deposited in the PDB, AlphaFold is expected to produce models analogous to the PDB structures. Since most of the benchmark targets in DB5.5 were included in AlphaFold training, there would be training bias associated with their predictions (i.e. our measured success rates are an upper bound). However, since both unbound and bound structures exist for the benchmark targets in the PDB, we first investigated whether AFm exhibits any bias towards either unbound or bound forms for the same protein sequence. Supplementary Fig.S1 compares the C*α*-RMSD of all protein partners (calculated on a per-chain basis) of the AFm predicted complex structures from the bound (B) and unbound (U) crystal structures on a log-log scale (a few AFm predicted models were 20 Å apart from both bound and unbound structures). As evident from Supplementary Fig.S1A, the protein partners from the AFm top-ranked model deviate from both unbound and bound forms and skew more often towards the bound state. Antibody-antigen targets further demonstrate a similar trend, however with fewer targets predicted within sub-angstrom accuracy to the bound form (29.7% for Ab-Ag targets as opposed to 41% for DB5.5). We also calculated the TM-scores^26^ of the AFm predicted complex structures with respect to the bound and the unbound crystal structures (Supplementary Fig.S2). As TM-scores reflect a global comparison between structures and are less sensitive to local structural deviations, no strong conclusions could be derived. This is in agreement with our intuition that since both unbound and bound states of proteins will share a similar fold, and AlphaFold can predict structures with high TM-scores in most cases, gauging the conformational deviations with TM-scores would be inconclusive.

### AlphaFold pLDDT provides a predictive confidence measure for backbone flexibility

AlphaFold employs multiple sequence alignments with a multi-track attention-based architecture to predict three-dimensional structures of proteins and complexes. Further, for each structural prediction, it provides a residue-level confidence measure: the predicted local-distance difference test (pLDDT), estimating the agreement between predicted model to an experimental structure based on the C*α* LDDT test (*Methods*). Tunyasuvunakool *et al*. analyzed pLDDT confidence measures for the human proteome demonstrating the correlation between lower pLDDT scores with higher disordered regions in protein structures.^27^ Building on this observation, we evaluated whether there is a correlation between AlphaFold pLDDT confidence metric and the experimental metrics of conformational change between unbound and bound structures. In this regard, we compared the computational (AF-pLDDT) and experimental (per-residue RMSD and LDDT) metrics against each other.

As a reference, we first superimposed the unbound partners over the bound structures and calculated residue-wise C*α* deviations to determine the per-residue RMSD_*BU*_ values. LDDT_*BU*_ was measured by calculating the local distance differences in the unbound structure relative to the bound form. These metrics capture the extent of motion in the unbound-bound transitions for each of the protein targets. Next, we compared the per-residue pLDDT score from AFm predicted monomer models with the experimental metrics. Fig. 1A,B shows the results for two representative protein targets: kinase-associated phosphastase in complex with phospho-CDK2 (1FQ1^28^) and TGF-*β* receptor with FKBP12 domain (1B6C^29^). In both cases, pLDDT confidence scores correlate with the experimental measurements of binding: pLDDT decreases as LDDT_*BU*_ decreases and RMSD_*BU*_ increases. This is further illustrated with the AF2 predicted structures of the two targets superimposed over the bound structures (Fig. 1C). In regions of low confidence/pLDDT (highlighted in *red*), the prediction is inaccurate, but higher confidence/pLDDT regions (highlighted in *blue*) have high accuracy of prediction with the bound form. The results for the benchmark set (Supplementary Fig. S3-S4) show similar trends for most targets. The pLDDT, thus, can suggest protein residues that move upon binding.

**Fig. 1.**
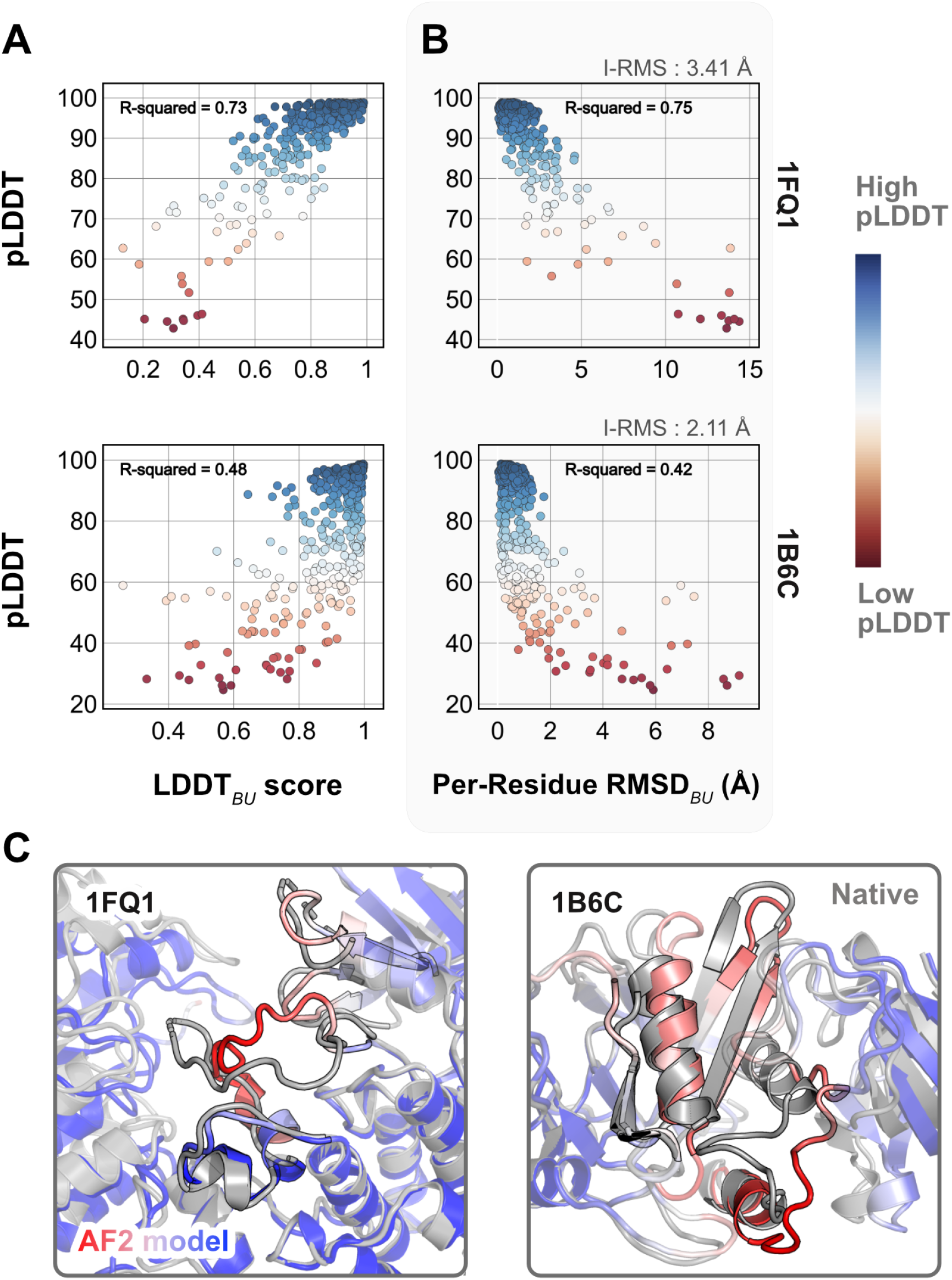
Comparison of AFm pLDDT with structural metrics. (A) AlphaFold pLDDT plotted against LDDT_*BU*_ (local distance difference test). LDDT_*BU*_ is calculated by comparing the unbound and bound environment for each residue. High scores correlate with high pLDDT (*red*). (B) Per-residue root-mean-square-deviation between unbound-bound structures (Per-Residue RMSD_*BU*_) v/s AlphaFold pLDDT for two example complex structures. Higher RMSDs correlate with lower pLDDT. (C) Structures for two targets (PDB ID: 1B6C and 1FQ1) with the experimental bound form (*in gray*) and the AlphaFold-multimer predicted model (*red-white-blue* in A and B). In both cases, the residues with low pLDDT scores (*red*) are the residues with incorrect conformation and more conformational change.

### Interface-pLDDT correlates with DockQ and discriminates poorly docked structures

When the prediction accuracy is lower, it is often evident from lower confidence metrics (such as average pLDDT or PAE). However, for AlphaFold-multimer complex predictions, the confidence metrics of the overall prediction do not correlate with the accuracy of the docked prediction, *i*.*e*., even if the complex exhibits higher confidence, the docking interfaces could be incorrect. Fig. 2 shows a few examples of failed AFm predictions including rigid (2FJU^30^), medium (5VNW^31^) and flexible targets (1IB1^32^, 2FJG^33^). In all the examples, the AFm model (highlighted in *red* to *blue* based on residue-wise pLDDT) is superimposed over an individual binding partner, and the bound structure is highlighted in *pale-green*. AFm models predict the individual subunits (protein partners) accurately in almost all scenarios, however the docking orientation is incorrect.

**Fig. 2.**
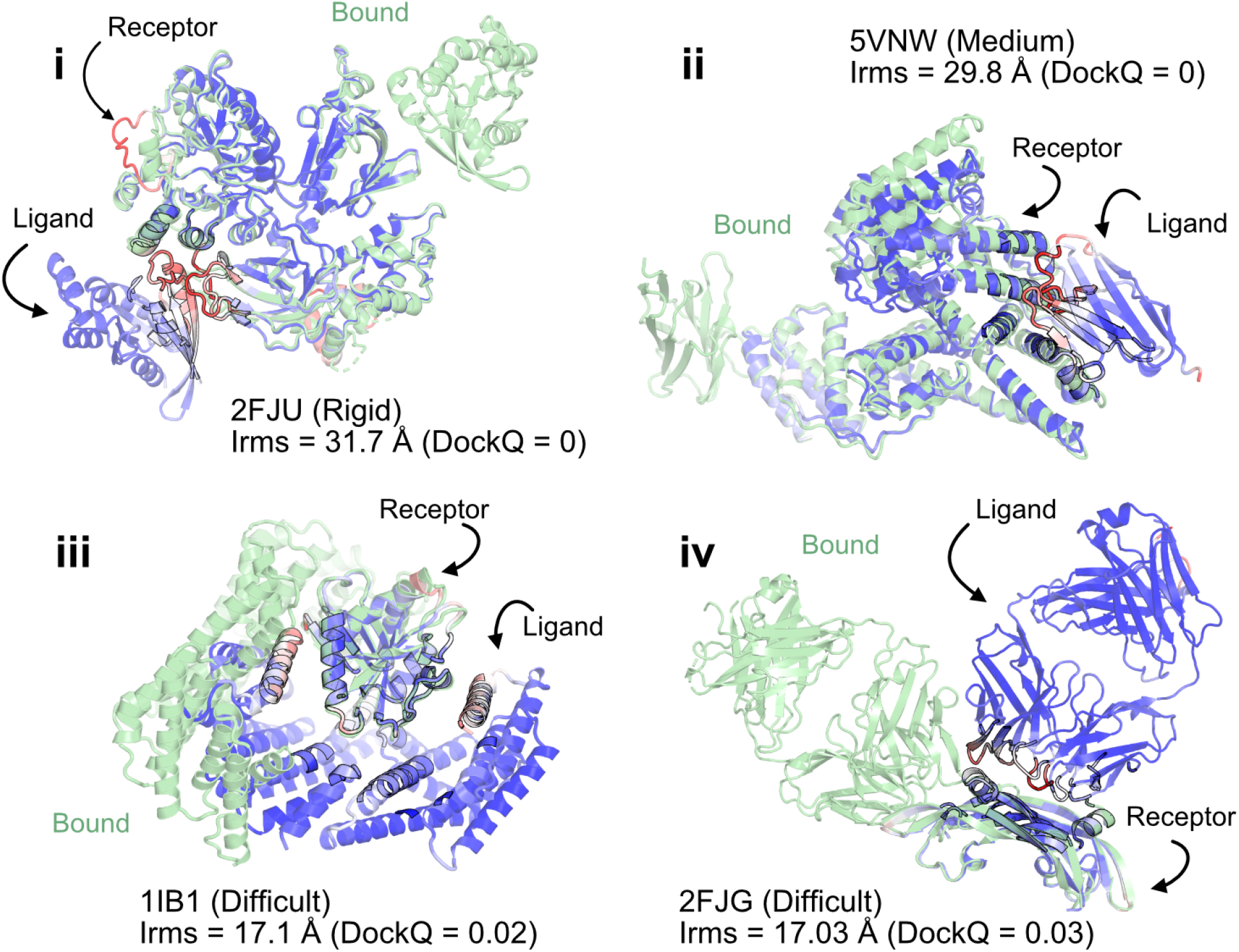
AlphaFold multimer predictions with reference to bound experimentally-characterized structures. Four targets with poor DockQ scores and high interface RMSDs: (i) activated Rac1 bound to phospholipase C*β*2 (2FJU) - rigid target (RMSD_UB_= 1.04 Å), (ii) nanobody bound to serum albumin (5VNW) - medium target (RMSD_UB_= 1.49 Å), (iii) 14-3-3 zeta Isoform:serotonin N-acetyltransferase complex (1IB1) - difficult target (RMSD_UB_= 2.09 Å), and (iv) G6 antibody in complex with the VEGF antigen - difficult target (RMSD_UB_= 2.51 Å). Bound structure in *green* and AlphaFold prediction colored by residue-wise pLDDT in *red* → *blue*. (low confidence → high confidence).

We investigated whether any of the AlphaFold predictive metrics could be repurposed for distinguishing native-like binding sites from non-native ones. That is, can one could utilize pLDDT or PAE from AFm models to determine whether the predicted docked complex has the accurate binding orientation? Thus, we evaluated accuracy with the DockQ score, the standard metric for docking model quality.^34^ DockQ ∈ [0, 1] combines interface RMSD (Irms), fraction of native-like contacts (*f*_nat_), and ligand-RMSD (Lrms). DockQ scores above 0.23 correspond to models with a CAPRI quality of “acceptable” or higher. As an acceptable quality target implies docked decoys are in the near-native binding region, we chose a binary classification of success with a threshold of DockQ = 0.23. We then tested how well DockQ correlated with several AFm-derived metrics: (a) Interface residues: the number of interface residues (atoms of residues on one partner within 8 Å from an atom on another partner); (b) Interface contacts: the number of interface contacts between the residues on the interface (C*β* atoms within 5 Å); (c) Average pLDDT, determined by averaging over the per-residue LDDT score of the entire protein complex; and (d) Interface-pLDDT, determined by averaging the per-residue LDDT score only over the predicted interfacial residues (as identified in case *a*).

Fig. 3A shows the classification accuracy of each of these metrics with a receiver-operating characteristics curve. The interface-pLDDT metric stands out with a higher true positive rate (TPR) with an area under curve (AUC) of 0.86. With interface-pLDDT as a discriminating metric, we tested multiple thresholds to estimate the optimum cut-off for distinguishing near-native structures (defined as an interface-RMSD < 4 Å) from the predictions. Fig. 3B summarizes the performance with a confusion matrix for the chosen interface-pLDDT cutoff of 85. 79% of the targets are classified accurately with a precision of 75%, thereby validating the utility of interface-pLDDT as a discriminating metric to rank the docking quality of the AFm complex structure predictions. With newer structure prediction tools such as AlphaFold3^35^ and ESM3^36^ being released, investigating features that could predict flexible residues or interface site would be valuable, as this information may guide local docking. This discrimination is also evident in the highlighted interface residues in Fig. 2, where the AFm predicted models have lower confidence at predicted interfaces (*red*). Finally, we show the trend between DockQ scores and interface-pLDDT for each target in Fig. 3C. The interface-pLDDT threshold of 85 (*dashed line*) thus can serve as the AlphaFold-derived metric to distinguish acceptable quality docked predictions from incorrect models.

**Fig. 3.**
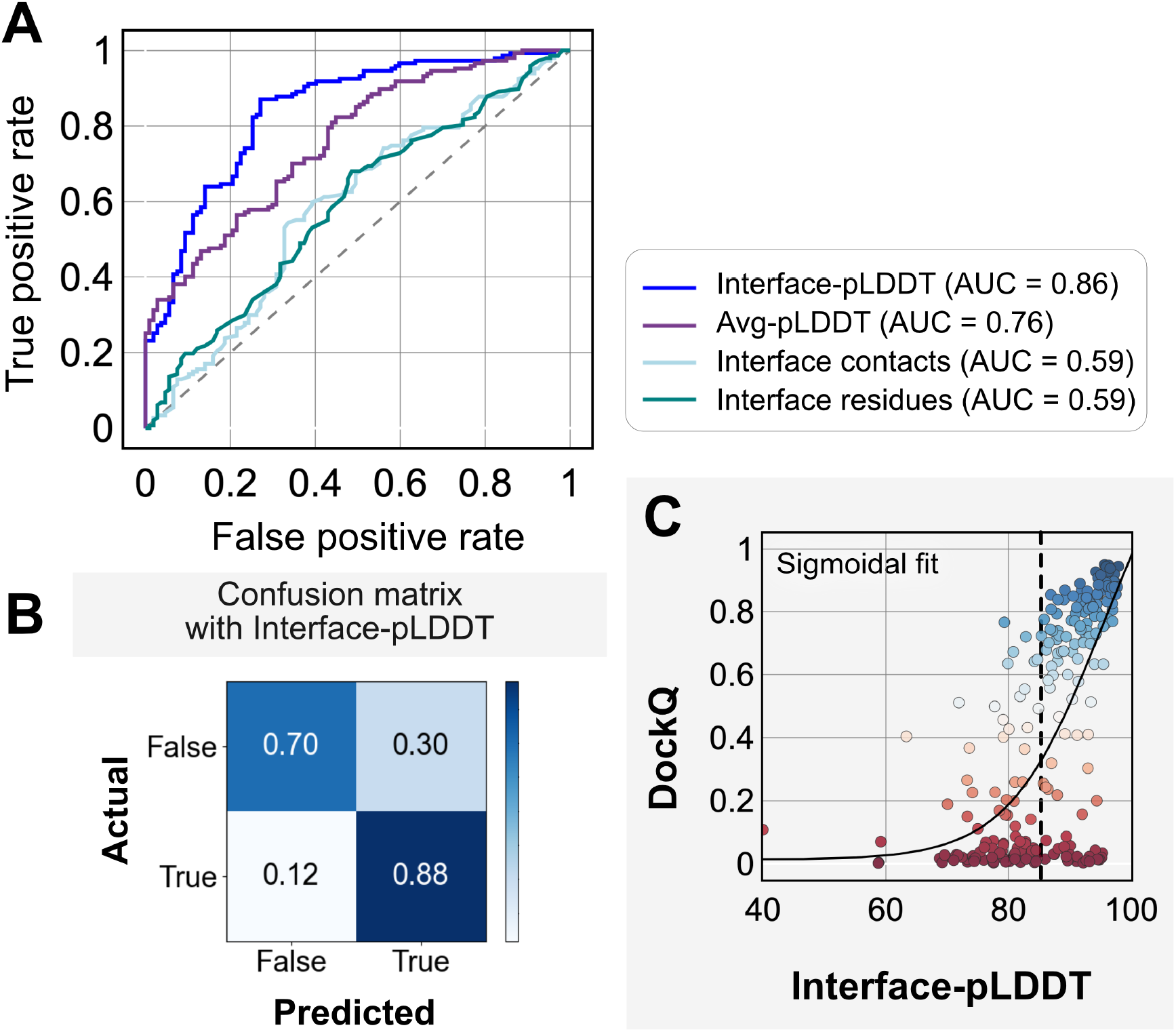
Interface-pLDDT is the best indicator of model docking quality. (A) Receiver-operator characteristics (ROC) curve as a function of different metrics for the docking dataset (n=254). Interface residues are defined based on whether atoms of residues on one partner are within 8 Å from atom/s on another partner. Interface-pLDDT is the average pLDDT of interface residues. Avg-pLDDT corresponds to the average pLDDT across all the residues in the predicted model. Interface contacts and interface residues are the counts of the interface contacts and interface residues respectively. Interface-pLDDT has the highest AUC score of 0.86. (B) Confusion matrix with an interface-pLDDT threshold between labels predicted false (<85) and true (≥85) and an interface-RMSD threshold between labels actually true (≤4 Å) and false(>4 Å) actual labels. (C) Interface-pLDDT versus DockQ for all protein targets in the benchmark set. DockQ is calculated from the predicted AlphaFold structure and the experimental bound structure in the PDB. We fit a sigmoidal curve to this available data.

### Docking benchmark targets initiated from AlphaFold models improves performance

With metrics to identify the flexible regions in the protein and the docking accuracy of generated docked models, we next fused AlphaFold-multimer (AFm) with our docking protocol, ReplicaDock 2.0^2^, to build a protocol for: (1) improving on incorrect AF docking predictions and producing alternate, near-native binding models and (2) capturing backbone conformational changes with our induced-fit protocol ReplicaDock2.0^2^. We named the protocol AlphaRED (AlphaFold-initiated Replica Exchange Docking). AlphaRED uses AFm predicted structures as the primary template, estimates docking accuracy metrics, and initiates global docking or refinement protocols as required.

Fig. 4 illustrates this docking pipeline. After AFm predicts a model from the protein sequences, we calculate the interface-pLDDT to determine the docking scheme to follow. If the AFm model is likely to be inaccurate (interface pLDDT < 85), we initiate a global replica exchange docking simulation to explore the protein conformational landscape and identify putative binding sites. On the other hand, if the interface-pLDDT > 85 for the AFm predicted model, the docked complex is likely in the correct binding orientation. This implies the global docking stage of the protocol can be skipped and local docking simulations can be directly initiated from the complex coordinates. Global docking follows an exhaustive, rigid-body search (no backbone moves) between the protein partners to sample putative landscapes in the energy landscape. An unbiased global docking simulation is initiated by randomizing the spatial orientation of protein partners from the input structure. The replica exchange MC routine ReplicaDock 2.0 performs rigid-body rotations (8°) and translations (4 Å). Sampled decoys are clustered from all replicas (based on energies and structural similarity) and the five top clusters are passed along for flexible local docking.

**Fig. 4.**
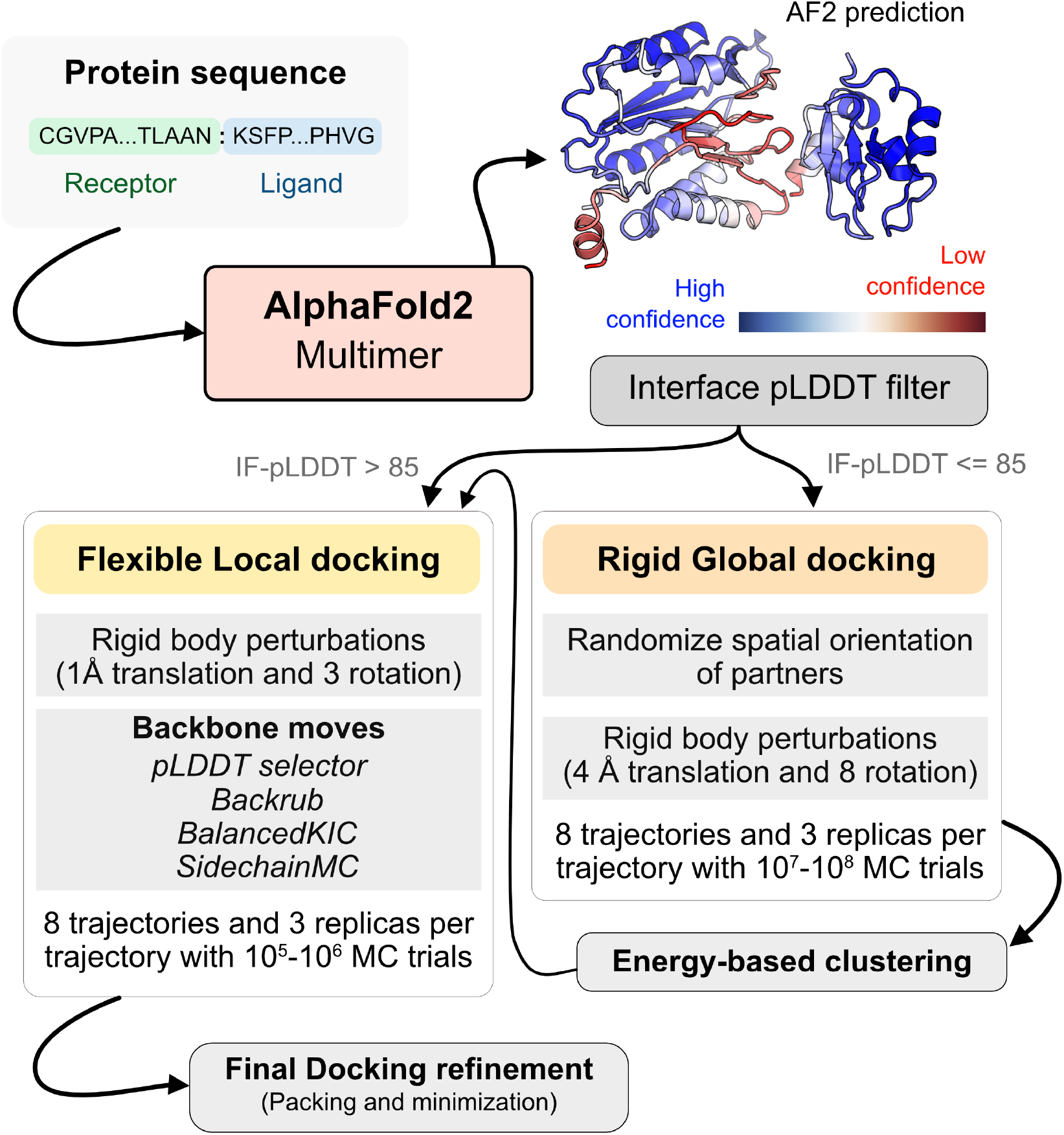
AlphaRED protein docking pipeline. Starting with protein sequences of putative complexes, we obtain predicted models from AlphaFold. Each model is accompanied with pLDDT scores, and based on the interface pLDDT we either initiate global rigid-body docking (interface pLDDT < 85), or flexible local docking refinement(interface pLDDT ≥ 85). For global rigid-body docking, the protein partners are first randomized in Cartesian coordinates and then docked with rigid-backbones using temperature replica exchange docking within ReplicaDock2.^2^ Decoy structures are clustered based on energy before flexible local docking refinement. In flexible local docking, we use the directed induced-fit strategy in ReplicaDock2. With mobile residues selected by the AlphaFold residue-wise pLDDT scores (threshold of 80). The protocol moves the backbones with Rosetta’s Backrub or Balanced Kinematic Closure movers. Output structures are refined and top-scoring structures are selected based on interface energy.

For flexible local docking, we perform aggressive backbone moves (backrub + kinematic closure, *Methods*) on candidate encounter complexes (clustered decoys), with fine rigid-body rotations and translations. To narrow conformational sampling, backbone moves are explicitly performed over residues identified as mobile’ based on the per-residue pLDDT metric (residue pLDDT < 80). Unlike ReplicaDock 2.0 that performs induced-fit over putative interfaces, this approach targets backbone motions over these predicted mobile residues, reducing the sampling space. Local docking decoys are further refined for side-chain packing and minimization to obtain the final docked structures (details in *Methods*). The methodological advancements and Rosetta movers in AlphaRED are further detailed in the *Methods* section.

We investigated AlphaRED’s performance on all 254 benchmark targets (Fig. 5). 97 targets under the threshold of interface-pLDDT (≤ 85) were passed to the global docking branch. Targets with interface-pLDDT over 85 proceeded directly to local docking refinement. For all benchmark targets, we compared AlphaRED performance of the top-scoring decoys against initial AFm-predicted complex structures. Fig. 5A shows the interface-RMSD (Irms) of the AFm and AlphaRED predictions from the bound structure, respectively. The lower Irms values indicate that AlphaRED improves on existing predictions for almost all targets. For targets where AFm prediction is determined to be a failure (interface-pLDDT ≤ 85, *red*), AlphaRED demonstrates a vast improvement in Irms for 93 out of 97 targets. Additionally, for targets where AFm prediction is considered acceptable (interface-pLDDT > 85), local docking slightly improves performance. AlphaRED captures lower interface-RMSDs (under 10 Å) for targets where AFm models dock at binding sites **∼**40 Å away. Fig. 5B demonstrates the improvement in recapitulating native-like contacts (*f*_nat_) with AlphaRED.

**Fig. 5.**
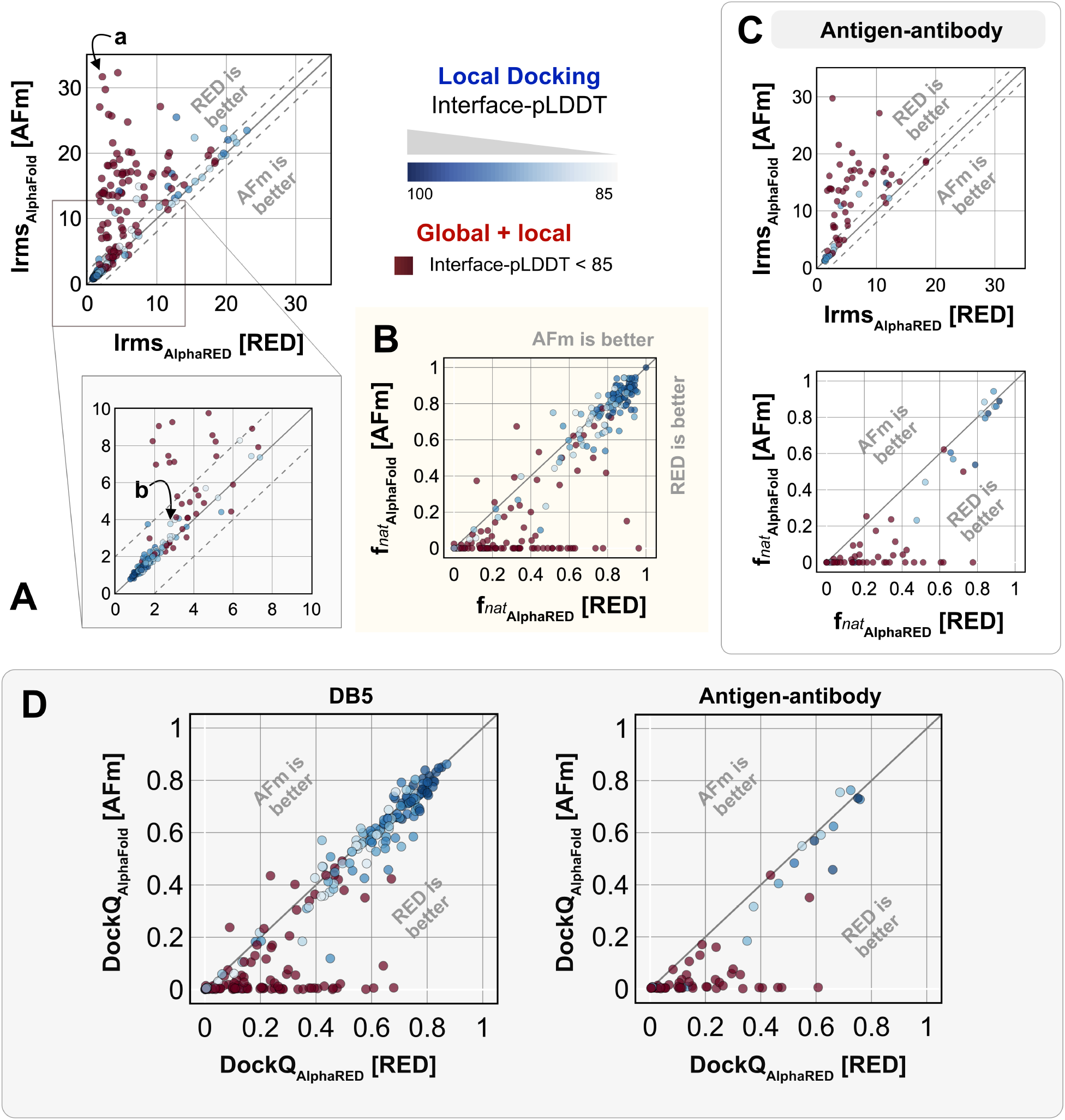
Docking performance. Targets with Interface-pLDDT ≤ 85 passed first to global rigid docking (*red*) where targets with interface-pLDDT> 85 proceeded directly to local flexible backbone docking refinement (colored based on their interface-pLDDT scores (in shades of *blue*) (A) Interface-RMSD from AlphaFold-multimer predicted models (*y-axis*) in comparison with AlphaRED models (*x-axis*). (B) Fraction of native-like contacts for models from AFm and AlphaRED respectively. (a) and (b) indicate two targets, (global and local docking) highlighted in Fig. 6. (C) Performance on the subset of antigen-antibody targets in DB5.5. (D) DockQ scores for the benchmark targets (DB5) and antibody-antigen complexes.

Fig. 5C shows the performance of the subset of antibody-antigen targets in the benchmark. Antibody targets are critical for understanding adaptive immune responses and for the design and engineering of antibody therapeutics.^37^ However, antibodies have proven challenging for deep learning methods, especially those reliant on multiple sequence alignments, as each antibody evolves in a different organism, and their antigens evolve on a different timescale altogether. Finally, to evaluate docking success rates, we calculate DockQ for top predictions from AFm and AlphaRED respectively (Fig. 5D). AlphaRED demonstrates a success rate (DockQ> 0.23) of 63% for the benchmark targets. Particularly for Ab-Ag complexes, AFm predicted acceptable or better quality docked structures in only 20% of the 67 targets. In contrast, the AlphaRED pipeline succeeds in 43% of the targets, a significant improvement. Most of the improvements in the success rates are for cases where AFm predictions are worse. For targets with good AFm predictions, AlphaRED refinement results in minimal improvements in docking accuracy.

Fig. 6 highlights a global docking (*a*) and local docking (*b*) example for targets 2FJU and 5C7X respectively. Starting from the incorrect AFm prediction (*orange*), AlphaRED samples over the conformational landscape to identify a top-scoring decoy (*blue*) with 2.6 Å Irms from the native (*gray*). Fig. 6b shows the extent of backbone sampling with ReplicaDock 2.0 local docking. The top-scoring decoy (*blue*) samples backbone closer to the bound form, improving model quality and docking accuracy.

**Fig. 6.**
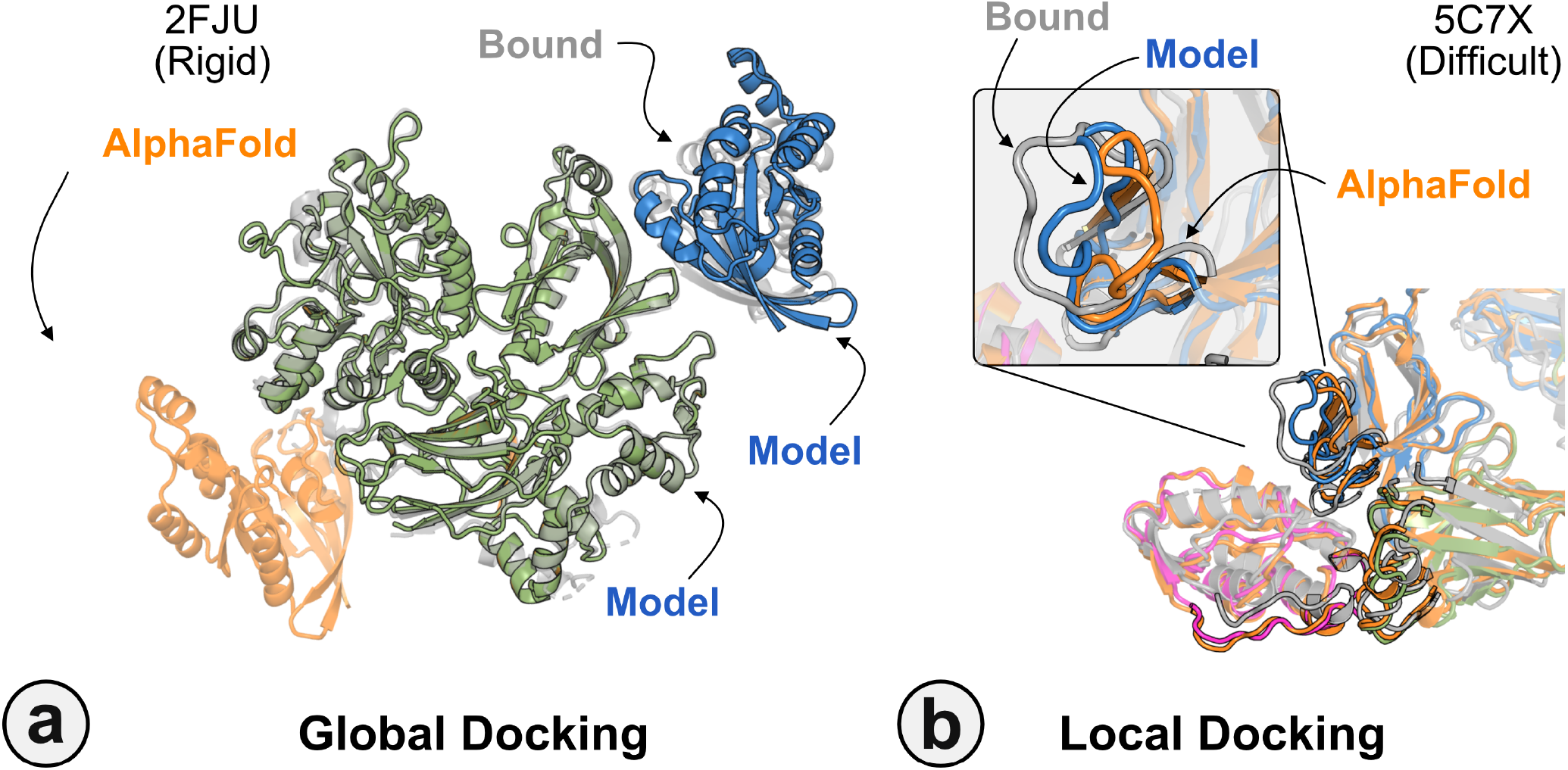
Global and local docking performance. Docking performance for targets (a) activated Rac1 bound to phos-pholipase C*β*2 (2FJU), and (b) neutralizing anti-human antibody Fab fragment in complex with human GM-CSF (5C7X). Starting from the AFm model (*orange*), global docking performance on 2FJU shows native-like binding site (*gray*) and sampled AlphaRED decoy (*blue*). For local docking, backbone sampling on mobile residues predicted by residue pLDDT (*outlined cartoon*) shows AlphaRED decoy (*blue*) moves backbone towards the bound form(*gray*).

### Evaluation on blind CASP15 targets

All results presented thus far may be biased by the fact that these benchmark target structures were used in the AFm training. The ultimate challenge for protein structure prediction protocols is to perform successfully over blind targets such as those in CASP (Critical Assessment of protein Structure Prediction) or CAPRI (Critical Assessment of PRotein Interactions) competitions.^8,38^ CASP15 (Summer 2022) provided multiple protein docking targets^24^ that were not included in AFm training, allowing an unbiased evaluation of our AlphaRED pipeline.^9^ Thus, we tested the protocol on the five heterodimeric nanobody-antigen complexes where most of the groups performed poorly (Fig. 7).

**Fig. 7.**
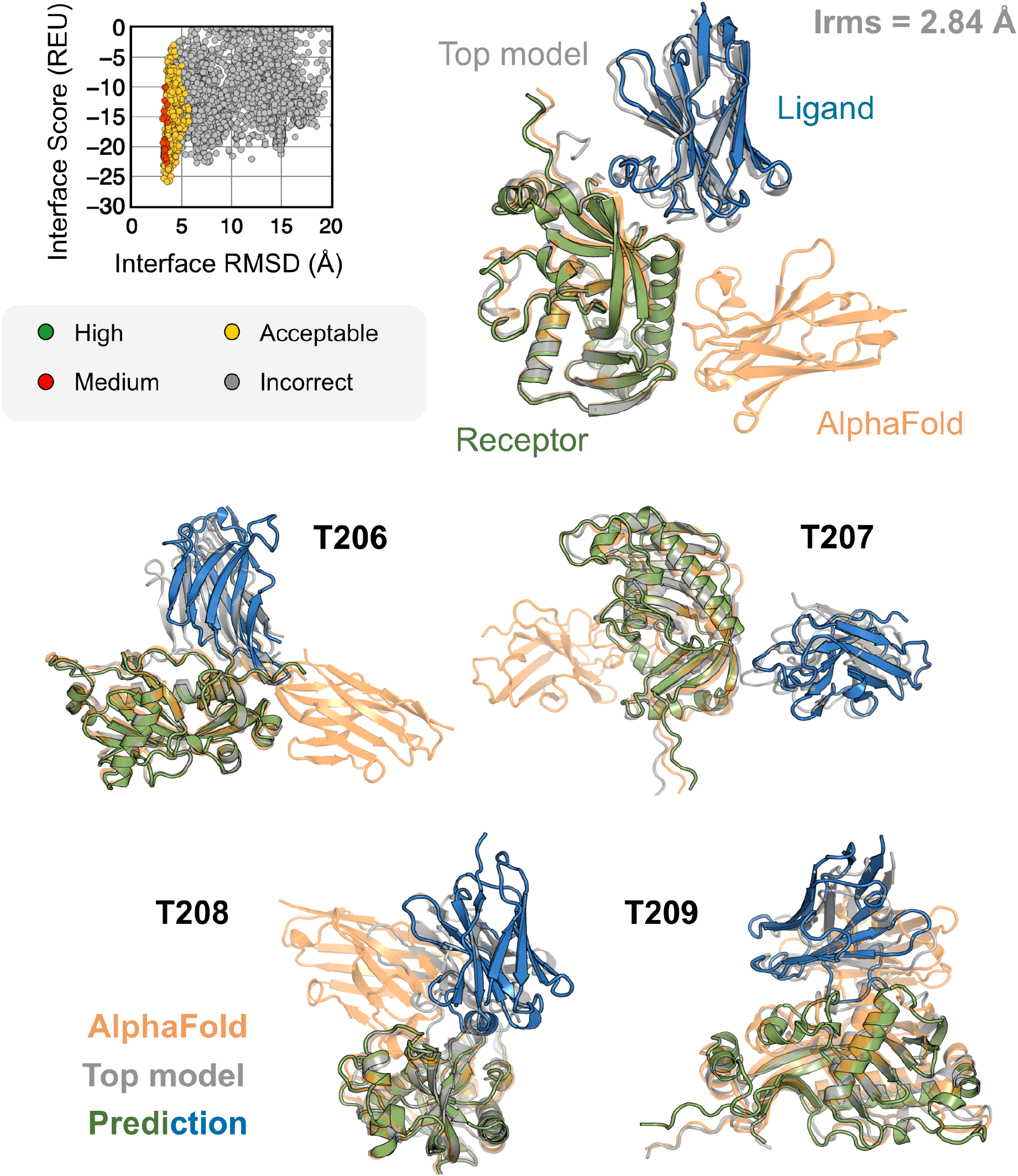
AFm and AlphaRED performance on CASP15 targets. Docking performance for CASP targets T205-T209. (*top*) T205. Interface score (Rosetta Energy Units, REU) vs Interface RMSD (Å) for candidate docking structures generated by the AlphaRED docking pipeline. (*top-right*) The top-scoring AlphaRED model (*green-blue*) recapitulates the native interface (*gray*) and has an interface RMSD of 2.84 Å. The distinction between the predicted model with respect to the AFm model (*orange*) is evident (*bottom*) Top-scoring AlphaRED predictions for targets T206, T207, T208, and T209 respectively.

Since the nanobody-antigen complexes were CASP targets, we did not have unbound structures, rather only the sequences of individual chains. Therefore, for each target, we employed the AlphaRED strategy as described in Fig. 4. All targets predicted with AFm had low interface-pLDDT thereby demanding global docking. This is unsurprising since the targets were nanobody-antigen targets and their CDRs, particularly CDR H3, are not conserved with a scarcity of co-evolution data with the antigen.^39^ For representative target T205, our docking strategy improves the performance drastically (interface RMSD 11.4 Å for AFm model to 2.84 Å for AlphaRED) and binds in the correct site. The interface scores versus interface-RMSD plot shows a distinct funnel with low-energy medium-quality structures (Fig. 6-*top*). Since the crystal structures are not yet released, the reference structure here is the top-model predicted for each category in CASP15.^9^ For all the targets, Fig. 6-*bottom* shows similar improvements for other nanobody-antigen complexes. These cases validate our strategy for blind targets, and demonstrate the ability of AlphaRED to serve as a robust pipeline, integrating AlphaFold with biophysical attributes to better predict protein complex structures.

## Discussion

AlphaFold has dramatically transformed the field of structural biology and is currently the state-of-the-art method to predict protein structures from sequences, not just for monomers but also for complexes and higher assemblies.^40^ One of the key elements of its success was the ability to mine evolutionary links between amino acids across protein families and determine structural templates. This approach dramatically improves prediction accuracy for monomers as reflected from prior CASP rounds. However, across protein interfaces, the evolutionary constraints can be weak and often skew predictions to inaccurate binding sites. Here we demonstrated how augmenting the predictions of AlphaFold with an energy-function dependent sampling approach reveals better backbone conformational diversity and accurate prediction of protein complex structures. By utilizing the AlphaRED strategy, we show that failure cases in AFm predicted models are improved for all targets (lower Irms for 97 of 254 failed targets) with CAPRI acceptable-quality or better models generated for 62% of targets overall (Fig. 8).

**Fig. 8.**
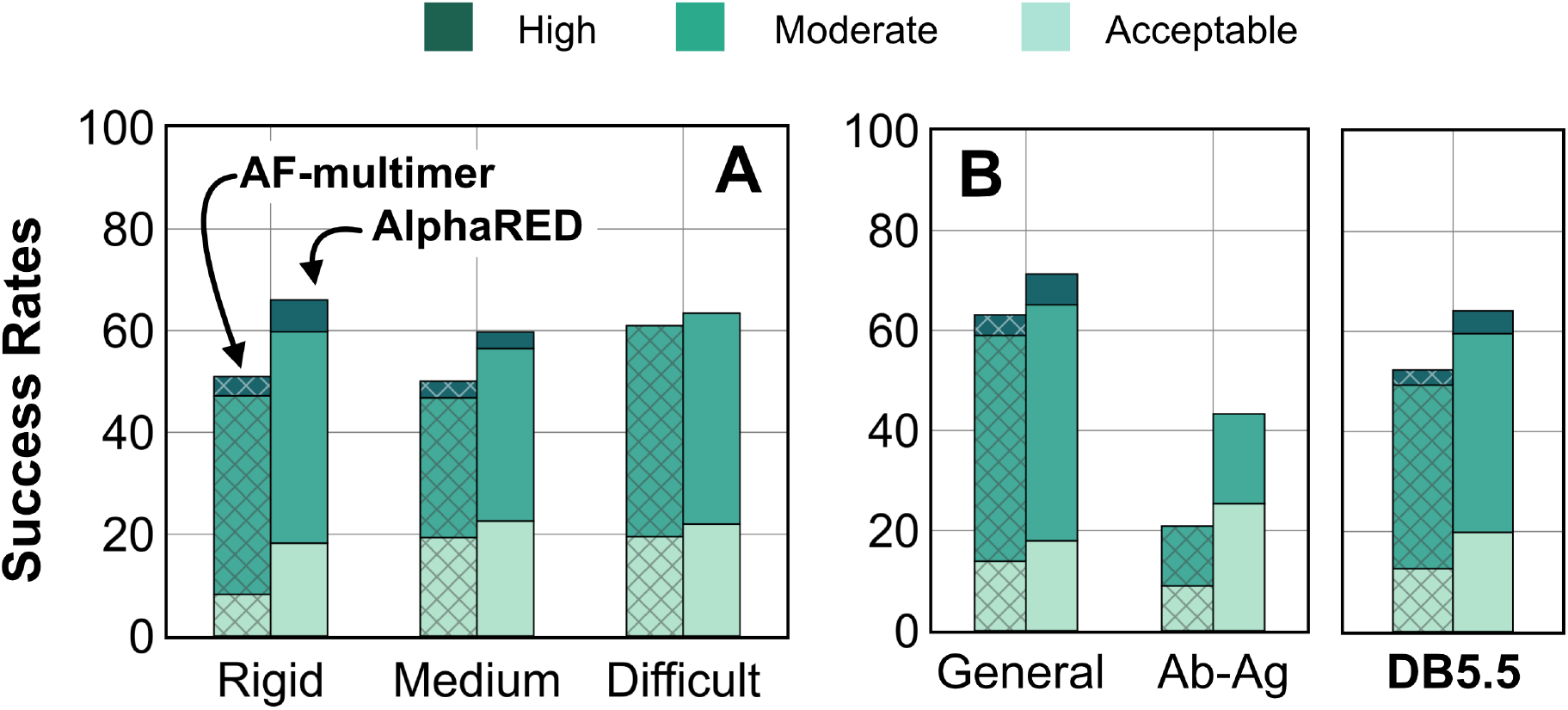
Docking prediction success with AFm and AlphaRED. Comparison of AFm (hashed) and AlphaRED performance for DB5.5 benchmark set. Success rates evaluated based on DockQ criteria: *incorrect* : DockQ < 0.23; *acceptable*: DockQ ∈ (0.23, 0.49]; *medium*: DockQ ∈ (0.49, 0.8]; *high*: DockQ ≥ 0.8 (A) Classification based on the scale of flexibility: difficult (35 targets); medium (60 targets); rigid (159 targets). (B) Performance on the antibody-antigen complexes (67 targets) and other (non-antibody targets).

First, we showed that AlphaFold confidence measures can be repurposed for estimating flexibility and docking accuracy. Interface-pLDDT, an average of the per-residue pLDDT only for the interfacial residues, is a robust metric to determine whether AFm predicted binding interfaces are correct. Additionally, thresholds of per-residue pLDDT can ascertain regions of backbone flexibility upon binding. Thus, AFm predicted models can be used as input structures for ReplicaDock 2.0 guiding the choices of global or local sampling and identifying the mobile protein segments. With DL-methods for structure prediction and downstream sampling with a physics-based energy function, one can efficiently explore the protein energy landscape as demonstrated with AlphaRED’s performance on DB5.5. Finally, we evaluated recent CASP15 targets to investigate the extrapolation of this strategy over blind protein targets. CASP15 targets were absent from the training routine of AlphaFold and served as blind challenges to determine the efficacy of the protocol. With AlphaRED, we obtained DockQ scores over 0.23 for all five targets, with medium-quality models (DockQ > 0.49) for targets T205, T207, and T208 respectively. AFSample, a top-performing group in CASP15, employed stochastic perturbation with dropout and increased sampling to obtain medium and high-quality models for these targets. However, AFSample requires GPU simulations to produce **∼**240x models with compute time **∼**1000x more than the baseline AFm.^10^ On other hand, we utilized ColabFold^11^ to generate 1-5 structures for our docking routine with the baseline version. As opposed to a couple of days on GPU (each GPU node contains up to 48 cores) utilized by AFSample, our docking routine fused with ColabFold uses 5-7 hours on our CPU cluster (runs on 1 node, with 24 cores, approximating to **∼**100 hours of CPU-hours per target). The AlphaRED docking strategy demonstrates a new and better way to predict protein complex structures within feasible compute times.

This work is particularly impactful for its success rate on antibody-antigen targets. Deep learning promises accurate design and optimization of antibody therapeutics^37^, but a lack of fast and accurate docking methods for antibodies prevent high-throughput computational screening. Additionally, this work is impactful because by integrating a physics-based method for refinement, the pipeline can potentially handle post-translationally modified proteins or non-canonical residue types that are not defined in ML approaches like AF.

With this work, we have built upon the recent advances in structural biology to develop a robust tool for protein docking. We fused deep-learning tools with conventional physics-based sampling tools to develop a pipeline that extracts the best outcomes of each methodology; where deep-learning methods generate accurate, static structures, and physics-based sampling provides diversity and better discrimination. The protein conformational landscape is vast and deep-learning tools such as AlphaFold provide a snapshot of relevant local minima that can aid in narrowing down the degrees of freedom in sampling.^41^ With the paradigm shift in computational structural biology towards deep-learning approaches, integrating physics within these models has tremendous potential towards understanding protein dynamics, modulating protein-protein interactions, and downstream applications to protein design.

## Methods

### Prediction of structures

For each target in the DB5.5 dataset, we first extracted the corresponding FASTA sequence for the bound complex and then obtained AlphaFold predicted models with the ColabFold v1.5.2^42^ implementation of AlphaFold^3^ and AlphaFold-multimer (v.2.3.0)^4^. Each prediction run was performed without templates, with automatic alignments and the default number of recycles to generate five relaxed predictions. Each AlphaFold prediction includes a per-residue pLDDT (predicted LDDT) measurement^43^, a confidence measure in prediction accuracy, and predicted template alignment (pTM) score.^26^ The models were structurally compared with the unbound and bound structures (deposited in the PDB) for measuring flexibility, similarity and accuracy of docking prediction.

### Metrics for backbone flexibility: RMSD and LDDT

Structures of proteins deposited in the PDB^44^ provide a static representation of the native-state of the protein. However, structural diversity has been captured by experimental techniques to identify different states of a protein in diverse physiological or chemical states, e.g. catalysis^45^, transport^46^, and ligand binding^47^. For protein docking challenges in particular, conformational changes are binding-induced, leading to structural differences between unbound and bound structures of protein targets.

To measure the conformational change in protein structures, we calculated two metrics: C*α* root-mean-square-deviation (RMSD) and local distance difference test (LDDT)^43^. To get a detailed representation of the intrinsic motion of a protein, we calculated RMSDs at a residue-level, *i*.*e*., per-residue C*α* RMSD for each residue of a protein target. The sequences+structures of unbound and bound proteins were aligned and the RMSDs were calculated for the aligned residues. The total sequence lengths were also matched and lingering end-termini residues were trimmed to ensure structural and sequential similarity.

Local Distance Difference Test (LDDT) is a superimposition-free score that estimates local distance differences in a model relative to a reference structure.^43^ Unlike the Global Distance Test (GDT)^48^ score based on rigid-body superimposition, the LDDT score measures the conserved local interactions in the protein model to the reference. For every residue, it computes the distance between all pair of atoms *D*(*i, j*) in both the model and the reference structure (bound) within a threshold (defined as the inclusion radius, generally set to 10 Å). For each pairwise distance in both distance vectors, if the distance is within the threshold, the distance is considered conserved and the fraction of conserved distances is calculated. The final LDDT score is the average of this fraction for the tolerances of 0.5, 1, 2, and 4 Å.

For a protein structure with *N* number of residues, the overall LDDT score can be given as follows:

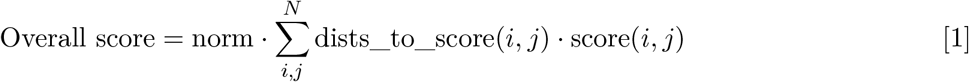

where norm is the normalization factor

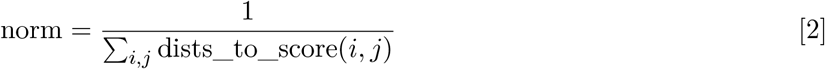

and score(*i, j*) is the LDDT score for the residue *i* with respect to every other residue *j*

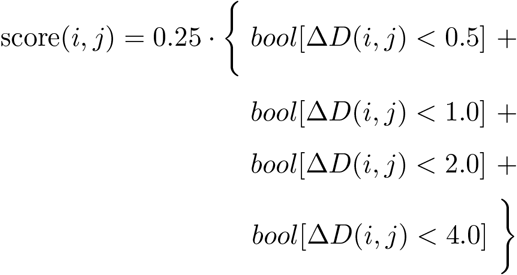

**Δ***D*(*i, j*) denotes the absolute difference between *D*_true_(*i, j*) and *D*_predicted_(*i, j*) calculated as follows:

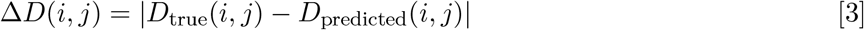

where *D*_true_(*i, j*) and *D*_predicted_(*i, j*) denote the distances between the C*α* coordinates of the *i*^*th*^ residue and the *j*^*th*^ residue for the true (reference) and predicted (model) structures respectively. 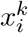 and 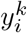 represent the *k*^*th*^ coordinate of the C*α* atom in the *i*^*th*^ residue in the reference (true) structure and predicted structure respectively, such that:

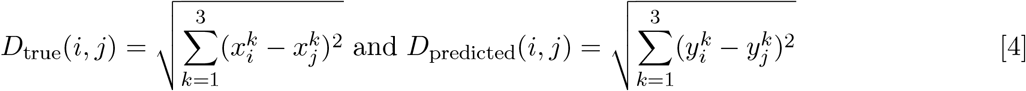

Finally, the distances to score (dists_to_score(*i, j*)) are computed as those pairwise distances within an inclusion radius (cutoff = 10 Å). 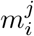 is the mask value (1 or 0) indicating if the *j*^*th*^ coordinate of the C*α* atom in the *i*^*th*^ residue exists in the true structure.

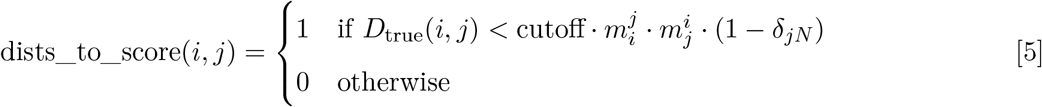

where *δ* = Kronecker Delta

The advantage of the LDDT measurement lies in the estimation of relative domain orientations in multi-domain proteins or concerted motions (e.g.: hinge-like moves in closed and apo proteins). In these cases, the RMSDs would be relatively high for all residues in the mobile domain, however, since the inter-residue distances within the domains are conserved, they would provide an inaccurate depiction of flexibility for the protein. Estimating both RMSDs and LDDT scores allows us to obtain a nuanced perspective of flexibility during protein association based on experimental structures.

### Developing a pipeline for protein docking

Using AlphaFold2 as a structural module, we built a pipeline for protein-protein docking to better predict protein complex structures with relatively higher accuracy. As illustrated in Fig. 4, given a sequence of a protein complex, we use the ColabFold implementation of AF2-multimer to obtain a predictive template. An interface-pLDDT filter determines the accuracy of the docking prediction of the top-ranked model from AFm. If the interface-pLDDT ≤ 85, the prediction has lower confidence in the docking orientation, and the protocol initiates a rigid, global docking search with ReplicaDock 2.0. Implementation of ReplicaDock 2.0 (global docking) is similar to the version reported in prior work^2^. Each simulation initiates 8 trajectories across 3 temperature replicas with inverse temperatures set to 1.5^-1^ kcal^−1^.mol, 3^−1^ kcal^−1^.mol and 5^−1^ kcal^−1^.mol, respectively. Across each replica within each trajectory, rigid body perturbations (4 Å translations and 8° rotations) are performed for an exhaustive global search. Next, we perform an energy-based clustering of the models to obtain diverse and energetically favourable clusters. Five cluster centers (decoys) are selected and passed to the flexible local docking stage to sample conformational changes.

On other hand, if the interface-pLDDT > 85, the binding orientation has higher confidence and the protocol directly performs a flexible local docking simulation skipping the rigid, global docking. In this stage, we perform smaller rigid-body perturbations (1 Å translations and 3° rotations) and aggressive backbone moves using a set of backbone and side-chain movers: Rosetta `Backrub`^49^, `Balanced Kinematic Closure` (BalancedKIC) and `Sidechain`. The sampling weights are biased such that backbone and side-chain movers are weighted higher than rigid body moves (3:1 weightage for backbone:rigid-body moves). We perform directed backbone sampling by focusing on predicted mobile residues (per residue pLDDT < 80). This is automated with the `BFactorResidueSelector` that selects contiguous sets of residues below the specified pLDDT threshold.

However, unlike the induced-fit strategy in ReplicaDock^2^, we perform backbone sampling directed only on the mobile residues (with per residue pLDDT < 80) identified from the AlphaFold model. We automate it using the `BFactorResidueSelector` to select contiguous sets of residues below the specified pLDDT threshold in the prior section. This residue subset is passed along to the backbone movers to sample backbone moves along with small rigid-body moves. Sampled decoyed are then refined, *i*.*e*. undergo side-chain packing and minimization, to output docked decoys. The best ranked decoys based on interface scores are then identified as the top-scoring structures.

## Data Availability

The source code for AlphaRED is available on github (github.com/Graylab/AlphaRED). An online server implementation is available on the Gray lab ROSIE server (rosie.graylab.jhu.edu). Users can submit their prediction and docking jobs on r2.graylab.jhu.edu/apps/submit/alpha-red.

## ACKNOWLEDGMENTS

The authors thank Sergey Ovchinnikov and Yoshitaka Moriwaki for ColabFold implementation of AlphaFold.

