## Supplementary File for "Reliable protein-protein docking with AlphaFold, Rosetta, and replica-exchange"

#### From sequences to structure to complexes: an *in-silico* pipeline for protein-protein docking

Figs. S1 to S4
Table S1
SI References

### Supplementary Methods

#### Benchmark Sets.

The benchmark set was curated from Docking Benchmark 5.5((1)), with targets classified based on their extent of flexibility, i.e., rigid, medium, and difficult. We also curated a subset of only antigen-antibody/nanobody targets from the overall set. For each target in the benchmark set, a FASTA sequence was obtained with individual chains separated by a colon (:) indicating chain break. This sequence was used for AlphaFold-multimer (AFm) structure prediction. For target 1N2C, AFm could not generate a structural prediction owing to the longer sequence length and is excluded from the benchmark. For each structural prediction that was generated, comparisons were made to its corresponding bound and unbound forms as obtained from the Docking Benchmark 5.5.

#### Metrics and evaluation.

The docking performance was evaluated based on interface RMSD (I-rms), fraction of native-like contacts ($f_{\texttt{nat}}$), CAPRI quality, and DockQ scores. These metrics are defined as follows:

Interface RMSD (I-rms): The root-mean-sqaure-deviation (RMSD) of all atoms on the interface in a docked protein structure relative to a reference structure (native). Interface residues are defined as all amino acid residues within 10 Å of any residue on the binding partner.

fraction of native-like contacts ($f_{\texttt{nat}}$): The fraction of native-like contacts recovered in the docked structure relative to the reference structure (native).

CAPRI quality: A CAPRI-based rank calculated on the basis of I-rms, $f_{\texttt{nat}}$, and ligand-RMSD to classify a docked model as incorrect, acceptable, medium, or high-quality prediction.

DockQ Score: Similar to CAPRI-quality, the DockQ scores estimates a score ($\in [0, 1]$) estimating the accuracy of the docked complexes. We calculated this score based on the methodology described in Basu *et al.*(2)

Interface score (Isc): The interface score is analogous to thermodynamic binding energy of protein association. This score is estimated by calculating the total score (gibbs free energy) of protein complex and then by subtracting individual (monomeric) scores of protein partners in absence of its partner. Mathematically, for proteins A and B forming a complex AB, it can be defined as follows:

$$\Delta\Delta G_{\text{interface}} = \Delta G_{\text{AB}} - \Delta G_{\text{A}} - \Delta G_{\text{B}}$$

### Supplementary Results

#### Data availability.

AlphaRED utilizes ColabFold for structure prediction with Rosetta-based docking. The source code for AlphaRED is available on github (github.com/Graylab/AlphaRED). To ensure ease of availabilty for researchers, we have implemented an online server on the Gray Lab ROSIE server (rosie.graylab.jhu.edu). The server would implement the AlphaRED pipeline and for input sequences would provide docked models with Rosetta energies for further analysis. We expect this implementation to be a great resource for modeling and better understanding protein-protein interactions.

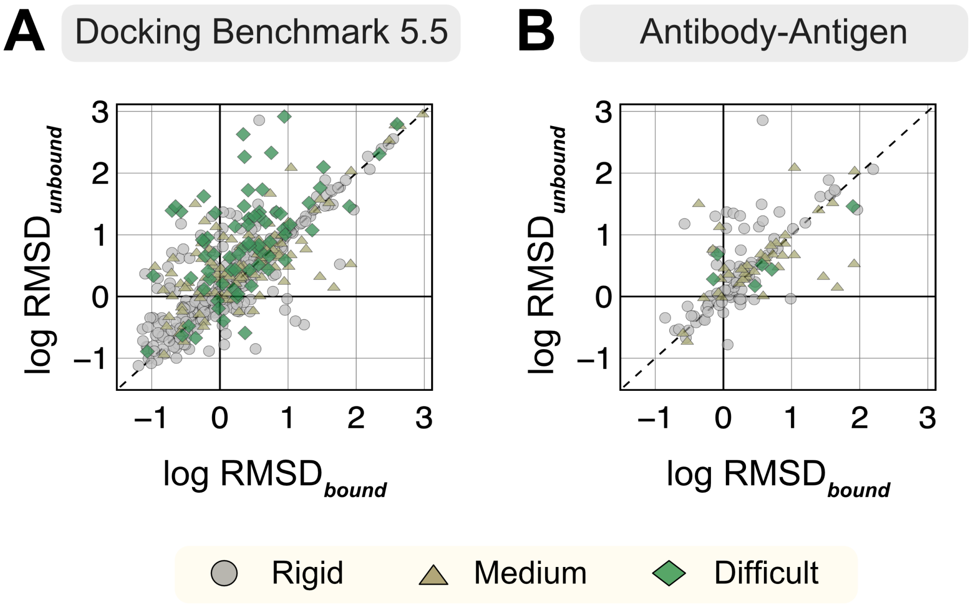


**Fig. S1. RMSDs of AlphaFold-multimer structures from experimental unbound and bound structures.** Distribution of the RMSD between the AlphaFold-multimer prediction top-ranked model and the experimental unbound and bound structures. For each target, the protein partners are split into receptor and ligand respectively for comparison. Each symbol represents a category of flexibility (rigid, medium, and flexible). (A) Dockground Benchmark set 5.5; (B) Antibody/nanobody-antigen targets from the benchmark.

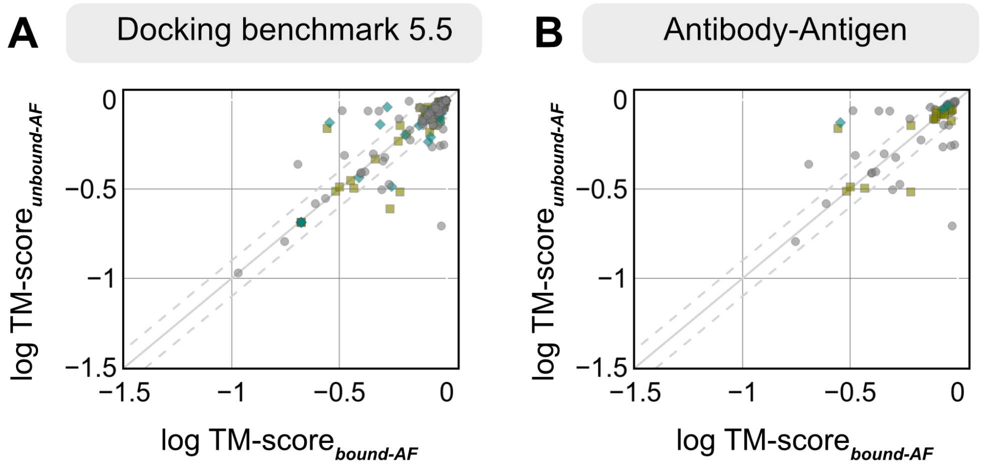


**Fig. S2. TM-scores of AlphaFold-multimer structures from experimental unbound and bound structures.** Distribution of the TM-score between the AlphaFold-multimer prediction top-ranked model and the experimental unbound and bound structures. For each target, the protein partners are split into receptor and ligand respectively for comparison. Each symbol represents a category of flexibility (rigid, medium, and flexible). (A) Dockground Benchmark set 5.5; (B) Antibody/nanobody-antigen targets from the benchmark.

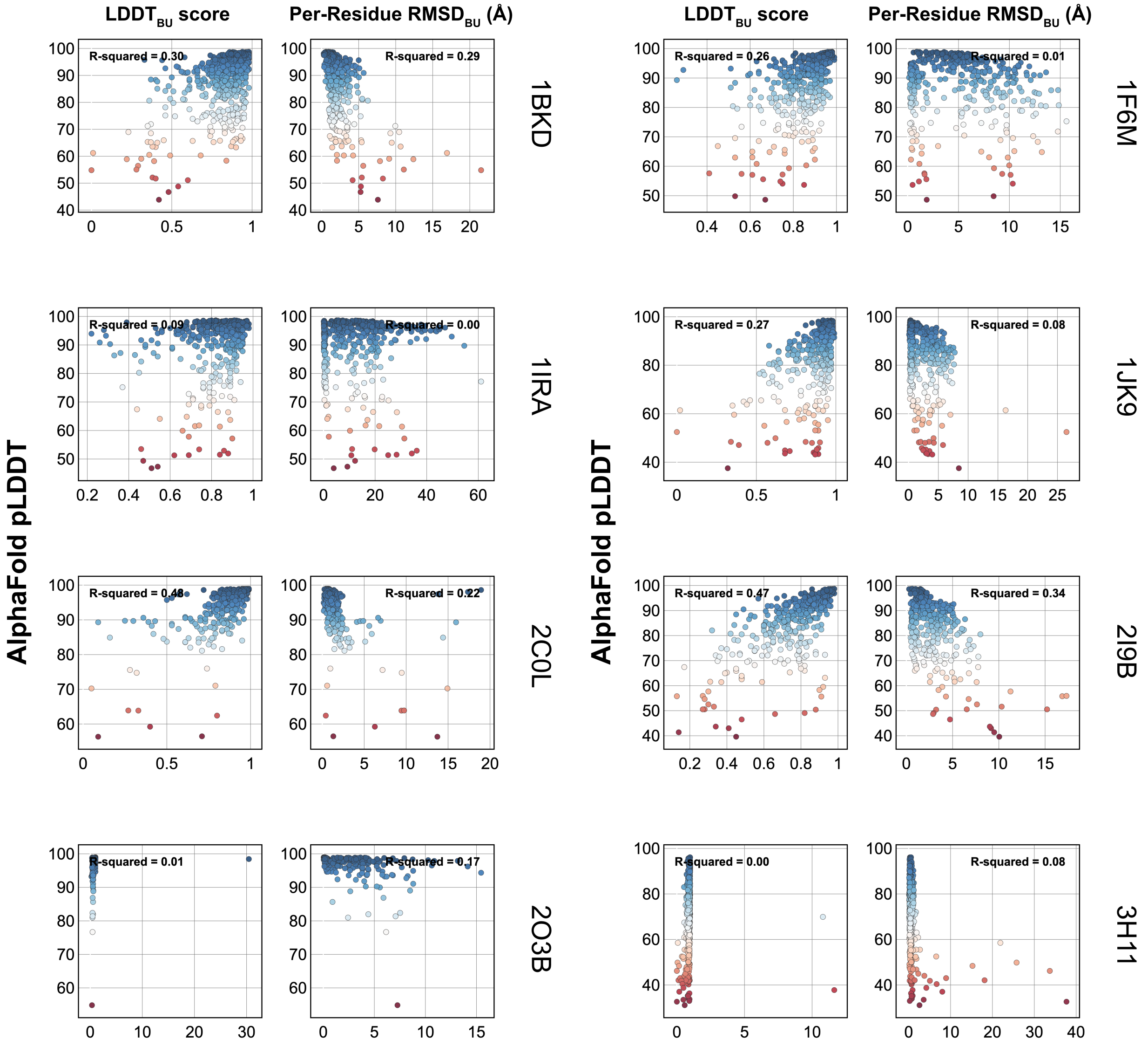


**Fig. S3. Comparison of AFm pLDDT with structural metrics.**

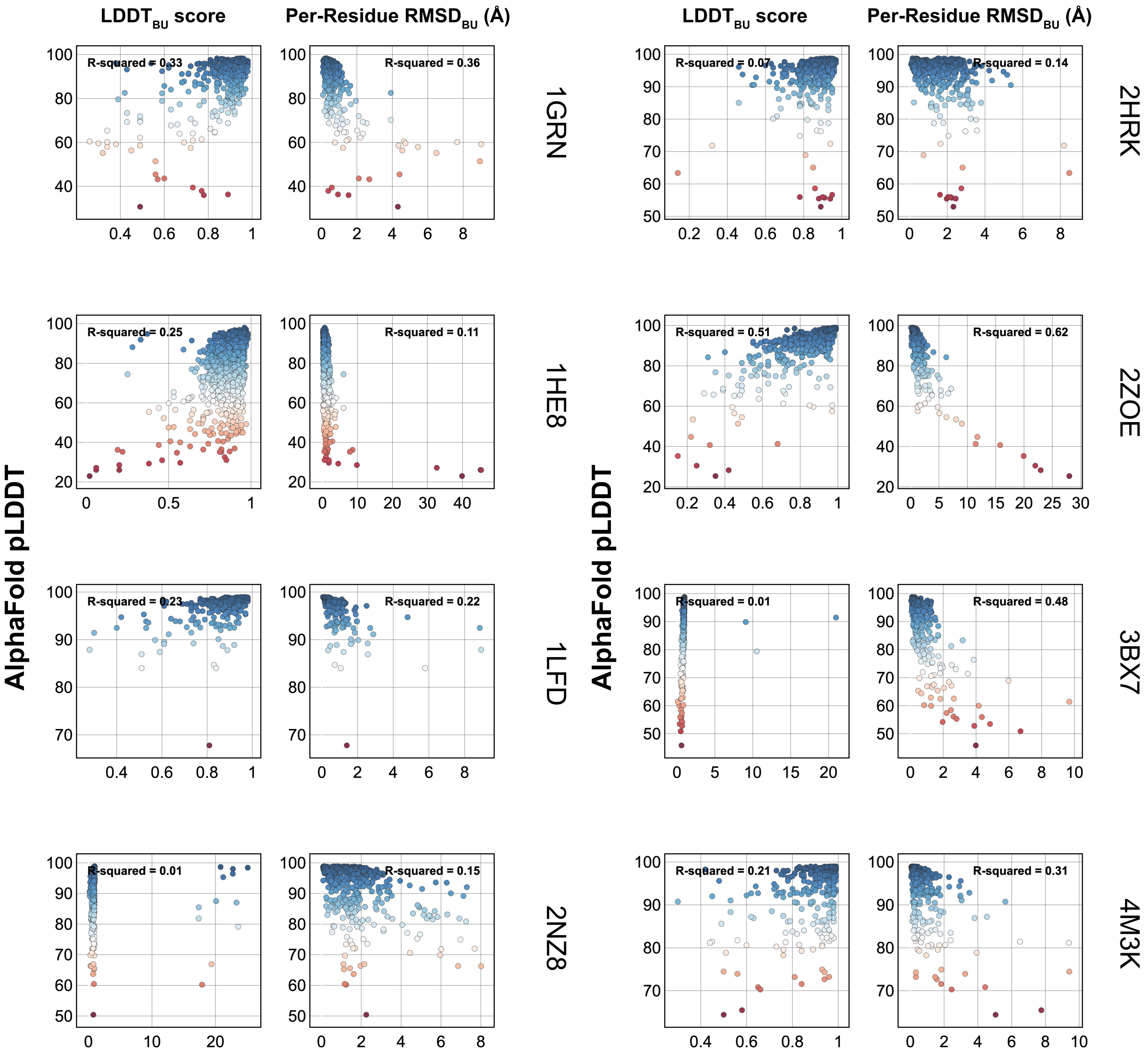


**Fig. S4. Comparison of AFm pLDDT with structural metrics.**

**Table S1. Performance of AlphaRED and AFm on Docking Benchmark Set 5.5**

| PDB | Interface RMSD | | $f_{nat}$ | | CAPRI Rank | | Flexibility |
|---|---|---|---|---|---|---|---|
| | AF2 | AlphaRED | AF2 | AlphaRED | AF2 | AlphaRED | |
| 1ACB | 2.51 | 2.02 | 0.86 | 0.85 | 1 | 1 | Difficult |
| 1ATN | 1.35 | 1.29 | 0.87 | 0.89 | 2 | 2 | Difficult |
| 1BKD | 1.64 | 1.48 | 0.84 | 0.83 | 2 | 2 | Difficult |
| 1DE4 | 4.43 | 5.91 | 0.37 | 0.12 | 0 | 0 | Difficult |
| 1E4K | 16.83 | 5.11 | 0.03 | 0.09 | 0 | 0 | Difficult |
| 1EER | 1.75 | 1.64 | 0.80 | 0.81 | 2 | 2 | Difficult |
| 1F6M | 2.76 | 2.75 | 0.50 | 0.56 | 1 | 1 | Difficult |
| 1FAK | 1.71 | 1.91 | 0.71 | 0.71 | 2 | 2 | Difficult |
| 1FQ1 | 4.06 | 3.24 | 0.50 | 0.56 | 0 | 1 | Difficult |
| 1H1V | 3.61 | 3.63 | 0.22 | 0.22 | 1 | 1 | Difficult |
| 1IBR | 1.34 | 1.31 | 0.79 | 0.89 | 2 | 2 | Difficult |
| 1IRA | 2.76 | 2.42 | 0.52 | 0.62 | 1 | 1 | Difficult |
| 1JK9 | 1.70 | 1.64 | 0.79 | 0.87 | 2 | 2 | Difficult |
| 1JMO | 4.10 | 3.69 | 0.72 | 0.82 | 0 | 1 | Difficult |
| 1JZD | 17.62 | 17.60 | 0.00 | 0.00 | 0 | 0 | Difficult |
| 1PXV | 1.30 | 1.37 | 0.89 | 0.91 | 2 | 2 | Difficult |
| 1R8S | 2.12 | 2.06 | 0.74 | 0.75 | 1 | 1 | Difficult |
| 1RKE | 4.31 | 4.12 | 0.57 | 0.64 | 0 | 0 | Difficult |
| 1Y64 | 7.44 | 6.93 | 0.40 | 0.35 | 0 | 0 | Difficult |
| 1ZLI | 1.84 | 1.79 | 0.86 | 0.88 | 2 | 2 | Difficult |
| 2C0L | 1.36 | 1.21 | 0.89 | 0.90 | 2 | 2 | Difficult |
| 2FJG | 17.03 | 5.65 | 0.02 | 0.07 | 0 | 0 | Difficult |
| 2I9B | 3.98 | 3.03 | 0.59 | 0.66 | 1 | 1 | Difficult |
| 2J7P | 2.74 | 2.76 | 0.38 | 0.43 | 1 | 1 | Difficult |
| 2O3B | 1.27 | 1.29 | 0.73 | 0.81 | 2 | 2 | Difficult |
| 2O3B | 1.16 | 1.16 | 0.81 | 0.85 | 2 | 2 | Difficult |
| 2OT3 | 1.97 | 2.01 | 0.69 | 0.70 | 2 | 1 | Difficult |
| 3AAD | 22.37 | 21.03 | 0.00 | 0.00 | 0 | 0 | Difficult |
| 3AAD | 21.60 | 21.48 | 0.00 | 0.00 | 0 | 0 | Difficult |
| 3F1P | 16.89 | 12.14 | 0.00 | 0.00 | 0 | 0 | Difficult |
| 3FN1 | 1.93 | 1.94 | 0.75 | 0.77 | 2 | 2 | Difficult |
| 3H11 | 5.89 | 6.00 | 0.35 | 0.55 | 0 | 0 | Difficult |
| 4DW2 | 18.37 | 7.00 | 0.00 | 0.02 | 0 | 0 | Difficult |
| 4GAM | 28.34 | 28.11 | 0.00 | 0.00 | 0 | 0 | Difficult |
| 5C7X | 3.76 | 2.80 | 0.23 | 0.48 | 1 | 1 | Difficult |
| 1B6C | 5.30 | 4.24 | 0.39 | 0.31 | 0 | 0 | Medium |
| 1CGI | 1.66 | 1.63 | 0.86 | 0.86 | 2 | 2 | Medium |

| PDB | Interface RMSD | | $f_{nat}$ | | CAPRI Rank | | Flexibility |
|---|---|---|---|---|---|---|---|
| | AF2 | AlphaRED | AF2 | AlphaRED | AF2 | AlphaRED | |
| **1FC2** | 1.96 | 1.87 | 0.84 | 0.88 | 2 | 2 | Medium |
| **1GP2** | 2.33 | 1.93 | 0.48 | 0.55 | 1 | 2 | Medium |
| **1GRN** | 1.80 | 1.72 | 0.78 | 0.76 | 2 | 2 | Medium |
| **1HE8** | 3.48 | 3.42 | 0.77 | 0.77 | 1 | 1 | Medium |
| **1I2M** | 3.03 | 3.10 | 0.74 | 0.66 | 1 | 1 | Medium |
| **1IB1** | 17.08 | 5.07 | 0.00 | 0.22 | 0 | 0 | Medium |
| **1IJK** | 5.18 | 5.23 | 0.78 | 0.72 | 0 | 0 | Medium |
| **1JIW** | 1.53 | 1.41 | 0.63 | 0.75 | 2 | 2 | Medium |
| **1K5D** | 1.55 | 1.57 | 0.75 | 0.75 | 2 | 2 | Medium |
| **1KKL** | 20.46 | 17.77 | 0.02 | 0.00 | 0 | 0 | Medium |
| **1LFD** | 1.00 | 1.07 | 0.88 | 0.94 | 2 | 2 | Medium |
| **1M10** | 4.40 | 5.40 | 0.27 | 0.33 | 0 | 0 | Medium |
| **1MQ8** | 1.97 | 1.58 | 0.73 | 0.94 | 2 | 2 | Medium |
| **1NW9** | 13.89 | 4.93 | 0.00 | 0.34 | 0 | 0 | Medium |
| **1R6Q** | 1.06 | 1.02 | 0.94 | 0.89 | 2 | 2 | Medium |
| **1SYX** | 1.22 | 1.17 | 0.79 | 0.90 | 2 | 2 | Medium |
| **1WQ1** | 1.54 | 1.45 | 0.68 | 0.79 | 2 | 2 | Medium |
| **1XQS** | 1.91 | 1.71 | 0.75 | 0.87 | 2 | 2 | Medium |
| **1ZM4** | 2.59 | 2.56 | 0.63 | 0.68 | 1 | 1 | Medium |
| **2CFH** | 3.09 | 2.89 | 0.75 | 0.63 | 0 | 1 | Medium |
| **2DD8** | 13.56 | 1.70 | 0.00 | 0.78 | 0 | 2 | Medium |
| **2H7V** | 2.26 | 2.55 | 0.68 | 0.73 | 1 | 1 | Medium |
| **2HRK** | 0.99 | 0.96 | 0.94 | 0.91 | 3 | 3 | Medium |
| **2NZ8** | 1.14 | 1.02 | 0.86 | 0.92 | 2 | 2 | Medium |
| **2OZA** | 13.24 | 8.20 | 0.00 | 0.17 | 0 | 0 | Medium |
| **2Z0E** | 1.25 | 1.37 | 0.93 | 0.93 | 2 | 2 | Medium |
| **3AAA** | 1.33 | 1.32 | 0.77 | 0.94 | 2 | 2 | Medium |
| **3AAD** | 22.37 | 15.42 | 0.00 | 0.00 | 0 | 0 | Medium |
| **3BX7** | 2.11 | 2.08 | 0.63 | 0.65 | 1 | 1 | Medium |
| **3CPH** | 5.63 | 4.09 | 0.14 | 0.21 | 0 | 0 | Medium |
| **3DAW** | 1.31 | 1.26 | 0.84 | 0.84 | 2 | 2 | Medium |
| **3EO1** | 21.58 | 3.63 | 0.00 | 0.33 | 0 | 1 | Medium |
| **3G6D** | 15.14 | 13.94 | 0.00 | 0.00 | 0 | 0 | Medium |
| **3HI6** | 4.89 | 4.52 | 0.17 | 0.26 | 0 | 0 | Medium |
| **3L5W** | 14.90 | 11.63 | 0.00 | 0.00 | 0 | 0 | Medium |
| **3R9A** | 19.05 | 4.63 | 0.00 | 0.18 | 0 | 0 | Medium |
| **3RJQ** | 9.77 | 4.75 | 0.00 | 0.21 | 0 | 0 | Medium |

| PDB | Interface RMSD | | $f_{nat}$ | | CAPRI Rank | | Flexibility |
|---|---|---|---|---|---|---|---|
| | AF2 | AlphaRED | AF2 | AlphaRED | AF2 | AlphaRED | |
| **3S9D** | 1.73 | 1.36 | 0.42 | 0.79 | 2 | 2 | Medium |
| **3SZK** | 14.92 | 7.40 | 0.00 | 0.00 | 0 | 0 | Medium |
| **3V6Z** | 17.17 | 6.25 | 0.00 | 0.06 | 0 | 0 | Medium |
| **4ETQ** | 20.17 | 5.91 | 0.00 | 0.15 | 0 | 0 | Medium |
| **4FZA** | 2.36 | 2.08 | 0.80 | 0.85 | 1 | 1 | Medium |
| **4IZ7** | 1.88 | 1.94 | 0.72 | 0.75 | 2 | 2 | Medium |
| **4JCV** | 20.06 | 11.80 | 0.00 | 0.02 | 0 | 0 | Medium |
| **4LW4** | 3.03 | 3.63 | 0.67 | 0.33 | 1 | 1 | Medium |
| **4M3K** | 16.13 | 2.06 | 0.00 | 0.46 | 0 | 1 | Medium |
| **4POU** | 13.79 | 2.43 | 0.00 | 0.51 | 0 | 1 | Medium |
| **5CBA** | 2.43 | 2.42 | 0.82 | 0.82 | 1 | 1 | Medium |
| **5E5M** | 5.26 | 3.15 | 0.04 | 0.40 | 0 | 1 | Medium |
| **5HGG** | 4.85 | 3.38 | 0.25 | 0.21 | 0 | 1 | Medium |
| **5HYS** | 16.42 | 9.16 | 0.00 | 0.00 | 0 | 0 | Medium |
| **5KOV** | 16.97 | 7.21 | 0.00 | 0.19 | 0 | 0 | Medium |
| **5VNW** | 29.75 | 2.58 | 0.00 | 0.42 | 0 | 1 | Medium |
| **5WHK** | 2.66 | 2.55 | 0.57 | 0.67 | 0 | 1 | Medium |
| **6A0Z** | 13.99 | 4.87 | 0.10 | 0.18 | 0 | 0 | Medium |
| **6AL0** | 3.09 | 2.78 | 0.44 | 0.52 | 0 | 1 | Medium |
| **6EY6** | 14.81 | 12.61 | 0.14 | 0.15 | 0 | 0 | Medium |
| **1A2K** | 24.77 | 5.79 | 0.00 | 0.14 | 0 | 0 | Rigid |
| **1AHW** | 1.93 | 1.41 | 0.54 | 0.79 | 2 | 2 | Rigid |
| **1AK4** | 7.37 | 7.37 | 0.17 | 0.23 | 0 | 0 | Rigid |
| **1AKJ** | 1.65 | 1.78 | 0.75 | 0.72 | 2 | 2 | Rigid |
| **1AVX** | 1.42 | 1.65 | 0.94 | 0.92 | 2 | 2 | Rigid |
| **1AY7** | 0.78 | 0.77 | 0.91 | 0.91 | 3 | 3 | Rigid |
| **1AZS** | 2.90 | 2.80 | 0.66 | 0.66 | 1 | 1 | Rigid |
| **1BUH** | 1.38 | 1.30 | 0.77 | 0.74 | 2 | 2 | Rigid |
| **1BVN** | 1.44 | 1.13 | 0.65 | 0.86 | 2 | 2 | Rigid |
| **1CLV** | 8.57 | 4.95 | 0.07 | 0.20 | 0 | 0 | Rigid |
| **1D6R** | 1.07 | 1.09 | 0.90 | 0.84 | 2 | 2 | Rigid |
| **1DFJ** | 1.45 | 1.57 | 0.77 | 0.83 | 2 | 2 | Rigid |
| **1DQJ** | 7.43 | 5.15 | 0.03 | 0.15 | 0 | 0 | Rigid |
| **1E6E** | 1.97 | 1.89 | 0.85 | 0.80 | 2 | 2 | Rigid |
| **1E6J** | 14.57 | 12.09 | 0.00 | 0.03 | 0 | 0 | Rigid |
| **1E96** | 1.49 | 1.23 | 0.86 | 0.92 | 2 | 2 | Rigid |
| **1EAW** | 1.11 | 1.19 | 0.78 | 0.81 | 2 | 2 | Rigid |

| PDB | Interface RMSD | | $f_{nat}$ | | CAPRI Rank | | Flexibility |
|---|---|---|---|---|---|---|---|
| | AF2 | AlphaRED | AF2 | AlphaRED | AF2 | AlphaRED | |
| **1EFN** | 15.08 | 3.34 | 0.00 | 0.33 | 0 | 1 | Rigid |
| **1EWY** | 5.70 | 4.60 | 0.06 | 0.11 | 0 | 0 | Rigid |
| **1EXB** | 23.79 | 19.64 | 0.00 | 0.05 | 0 | 0 | Rigid |
| **1EZU** | 25.50 | 12.81 | 0.00 | 0.01 | 0 | 0 | Rigid |
| **1F34** | 1.94 | 1.94 | 0.73 | 0.71 | 2 | 2 | Rigid |
| **1F51** | 19.73 | 19.91 | 0.00 | 0.00 | 0 | 0 | Rigid |
| **1FCC** | 27.07 | 1.76 | 0.00 | 0.74 | 0 | 2 | Rigid |
| **1FFW** | 1.41 | 1.28 | 0.86 | 0.96 | 2 | 2 | Rigid |
| **1FLE** | 1.52 | 1.46 | 0.81 | 0.91 | 2 | 2 | Rigid |
| **1FQJ** | 2.20 | 2.30 | 0.74 | 0.76 | 1 | 1 | Rigid |
| **1GCQ** | 14.19 | 4.55 | 0.00 | 0.15 | 0 | 0 | Rigid |
| **1GHQ** | 13.10 | 12.54 | 0.00 | 0.00 | 0 | 0 | Rigid |
| **1GL1** | 0.80 | 0.94 | 0.90 | 0.87 | 3 | 3 | Rigid |
| **1GLA** | 4.12 | 3.10 | 0.22 | 0.30 | 0 | 1 | Rigid |
| **1GPW** | 1.45 | 1.38 | 0.89 | 0.87 | 2 | 2 | Rigid |
| **1GXD** | 25.92 | 3.05 | 0.00 | 0.51 | 0 | 1 | Rigid |
| **1H9D** | 1.57 | 1.46 | 0.84 | 0.89 | 2 | 2 | Rigid |
| **1HCF** | 14.46 | 14.56 | 0.00 | 0.00 | 0 | 0 | Rigid |
| **1HE1** | 1.80 | 1.73 | 0.78 | 0.82 | 2 | 2 | Rigid |
| **1HIA** | 1.19 | 1.23 | 0.87 | 0.95 | 2 | 2 | Rigid |
| **1I4D** | 20.11 | 6.39 | 0.00 | 0.16 | 0 | 0 | Rigid |
| **1J2J** | 1.21 | 1.18 | 0.88 | 0.92 | 2 | 2 | Rigid |
| **1JPS** | 16.85 | 11.19 | 0.02 | 0.04 | 0 | 0 | Rigid |
| **1JTD** | 1.15 | 0.94 | 0.75 | 0.83 | 2 | 3 | Rigid |
| **1JTG** | 2.29 | 2.10 | 0.60 | 0.60 | 1 | 1 | Rigid |
| **1JWH** | 1.78 | 1.63 | 0.86 | 0.83 | 2 | 2 | Rigid |
| **1K74** | 1.75 | 1.78 | 0.87 | 0.87 | 2 | 2 | Rigid |
| **1KAC** | 18.09 | 3.05 | 0.00 | 0.34 | 0 | 1 | Rigid |
| **1KLU** | 13.54 | 3.04 | 0.00 | 0.56 | 0 | 1 | Rigid |
| **1KTZ** | 1.06 | 1.06 | 0.89 | 0.93 | 2 | 2 | Rigid |
| **1KXP** | 1.17 | 1.17 | 0.85 | 0.83 | 2 | 2 | Rigid |
| **1M27** | 8.25 | 1.90 | 0.15 | 0.90 | 0 | 2 | Rigid |
| **1MAH** | 1.42 | 1.42 | 0.78 | 0.79 | 2 | 2 | Rigid |
| **1ML0** | 19.64 | 17.41 | 0.00 | 0.02 | 0 | 0 | Rigid |
| **1MLC** | 10.89 | 3.99 | 0.00 | 0.15 | 0 | 1 | Rigid |
| **1OC0** | 1.11 | 1.31 | 0.94 | 0.85 | 2 | 2 | Rigid |
| **1OFU** | 1.34 | 1.28 | 0.94 | 0.94 | 0 | 2 | Rigid |

| PDB | Interface RMSD | | $f_{nat}$ | | CAPRI Rank | | Flexibility |
|---|---|---|---|---|---|---|---|
| | AF2 | AlphaRED | AF2 | AlphaRED | AF2 | AlphaRED | |
| **1OPH** | 2.11 | 1.48 | 0.82 | 0.86 | 1 | 2 | Rigid |
| **1OYV** | 8.29 | 6.34 | 0.68 | 0.78 | 0 | 0 | Rigid |
| **1PPE** | 2.00 | 1.88 | 0.82 | 0.82 | 2 | 2 | Rigid |
| **1PVH** | 18.12 | 2.88 | 0.00 | 0.35 | 0 | 1 | Rigid |
| **1QA9** | 1.76 | 1.63 | 0.64 | 0.77 | 2 | 2 | Rigid |
| **1R0R** | 0.86 | 0.89 | 0.91 | 0.93 | 3 | 3 | Rigid |
| **1RLB** | 19.55 | 19.32 | 0.00 | 0.00 | 0 | 0 | Rigid |
| **1RV6** | 23.47 | 22.98 | 0.00 | 0.00 | 0 | 0 | Rigid |
| **1S1Q** | 14.25 | 2.42 | 0.00 | 0.49 | 0 | 1 | Rigid |
| **1S78** | 27.14 | 10.50 | 0.00 | 0.06 | 0 | 0 | Rigid |
| **1SBB** | 25.69 | 3.49 | 0.00 | 0.41 | 0 | 1 | Rigid |
| **1T6B** | 12.23 | 7.85 | 0.02 | 0.20 | 0 | 0 | Rigid |
| **1TMQ** | 12.66 | 12.73 | 0.05 | 0.08 | 0 | 0 | Rigid |
| **1UDI** | 1.23 | 1.16 | 0.77 | 0.92 | 2 | 2 | Rigid |
| **1US7** | 0.76 | 0.76 | 0.93 | 0.94 | 0 | 3 | Rigid |
| **1VFB** | 13.92 | 2.07 | 0.00 | 0.61 | 0 | 1 | Rigid |
| **1WDW** | 35.72 | 35.36 | 0.00 | 0.00 | 0 | 0 | Rigid |
| **1WEJ** | 16.95 | 9.41 | 0.00 | 0.07 | 0 | 0 | Rigid |
| **1XD3** | 1.72 | 1.71 | 0.94 | 0.89 | 2 | 2 | Rigid |
| **1XU1** | 13.59 | 13.07 | 0.00 | 0.00 | 0 | 0 | Rigid |
| **1YVB** | 1.14 | 1.12 | 0.83 | 0.87 | 2 | 2 | Rigid |
| **1Z0K** | 1.75 | 1.53 | 0.65 | 0.81 | 2 | 2 | Rigid |
| **1Z5Y** | 2.12 | 2.12 | 0.81 | 0.86 | 1 | 1 | Rigid |
| **1ZHH** | 16.31 | 16.53 | 0.00 | 0.00 | 0 | 0 | Rigid |
| **1ZHI** | 1.06 | 1.22 | 0.94 | 0.91 | 2 | 2 | Rigid |
| **2A1A** | 1.88 | 1.95 | 0.84 | 0.75 | 2 | 2 | Rigid |
| **2A5T** | 1.48 | 1.42 | 0.83 | 0.85 | 2 | 2 | Rigid |
| **2A9K** | 16.93 | 4.29 | 0.00 | 0.31 | 0 | 0 | Rigid |
| **2ABZ** | 8.93 | 6.97 | 0.08 | 0.10 | 0 | 0 | Rigid |
| **2AJF** | 16.70 | 5.00 | 0.00 | 0.22 | 0 | 0 | Rigid |
| **2AYO** | 2.10 | 2.11 | 0.68 | 0.57 | 1 | 1 | Rigid |
| **2B42** | 1.22 | 1.21 | 0.82 | 0.87 | 2 | 2 | Rigid |
| **2B4J** | 18.53 | 3.72 | 0.00 | 0.26 | 0 | 1 | Rigid |
| **2BTF** | 1.15 | 1.11 | 0.82 | 0.85 | 2 | 2 | Rigid |
| **2FD6** | 16.35 | 12.52 | 0.00 | 0.03 | 0 | 0 | Rigid |
| **2FJU** | 31.69 | 2.15 | 0.00 | 0.59 | 0 | 1 | Rigid |
| **2G77** | 1.87 | 2.16 | 0.75 | 0.52 | 2 | 1 | Rigid |

| PDB | Interface RMSD | | $f_{nat}$ | | CAPRI Rank | | Flexibility |
|---|---|---|---|---|---|---|---|
| | AF2 | AlphaRED | AF2 | AlphaRED | AF2 | AlphaRED | |
| **2GAF** | 2.45 | 2.94 | 0.53 | 0.44 | 1 | 1 | Rigid |
| **2GTP** | 1.00 | 0.96 | 0.79 | 0.90 | 3 | 3 | Rigid |
| **2HLE** | 2.18 | 1.92 | 0.64 | 0.79 | 1 | 2 | Rigid |
| **2HQS** | 2.49 | 2.38 | 0.56 | 0.60 | 1 | 1 | Rigid |
| **2I25** | 7.45 | 2.70 | 0.11 | 0.36 | 0 | 1 | Rigid |
| **2J0T** | 1.15 | 1.16 | 0.95 | 0.95 | 2 | 2 | Rigid |
| **2MTA** | 22.03 | 20.25 | 0.00 | 0.00 | 0 | 0 | Rigid |
| **2O8V** | 7.93 | 7.29 | 0.20 | 0.40 | 0 | 0 | Rigid |
| **2OOB** | 9.07 | 2.22 | 0.00 | 0.55 | 0 | 1 | Rigid |
| **2OOR** | 19.99 | 19.61 | 0.00 | 0.00 | 0 | 0 | Rigid |
| **2OUL** | 0.86 | 0.85 | 0.94 | 0.93 | 3 | 3 | Rigid |
| **2PCC** | 1.28 | 1.47 | 0.90 | 0.85 | 2 | 2 | Rigid |
| **2SIC** | 0.94 | 0.89 | 0.88 | 0.91 | 3 | 3 | Rigid |
| **2SNI** | 1.03 | 0.99 | 0.84 | 0.89 | 2 | 3 | Rigid |
| **2UUY** | 1.01 | 0.96 | 0.90 | 0.94 | 2 | 3 | Rigid |
| **2VDB** | 0.94 | 0.86 | 0.83 | 0.89 | 3 | 3 | Rigid |
| **2VIS** | 12.94 | 7.07 | 0.00 | 0.02 | 0 | 0 | Rigid |
| **2VXT** | 2.20 | 2.14 | 0.82 | 0.86 | 1 | 1 | Rigid |
| **2W9E** | 13.91 | 2.27 | 0.00 | 0.62 | 0 | 1 | Rigid |
| **2X9A** | 0.98 | 0.94 | 0.89 | 0.89 | 3 | 3 | Rigid |
| **2YVJ** | 3.74 | 1.65 | 0.10 | 0.45 | 1 | 2 | Rigid |
| **3A4S** | 1.64 | 1.68 | 0.85 | 0.90 | 2 | 2 | Rigid |
| **3BIW** | 11.77 | 10.32 | 0.10 | 0.10 | 0 | 0 | Rigid |
| **3BP8** | 32.31 | 4.38 | 0.00 | 0.44 | 0 | 0 | Rigid |
| **3D5S** | 1.85 | 1.67 | 0.73 | 0.76 | 2 | 2 | Rigid |
| **3EOA** | 17.43 | 11.54 | 0.00 | 0.09 | 0 | 0 | Rigid |
| **3H2V** | 6.98 | 2.07 | 0.00 | 0.63 | 0 | 1 | Rigid |
| **3HMX** | 19.54 | 11.61 | 0.00 | 0.03 | 0 | 0 | Rigid |
| **3K75** | 1.20 | 1.12 | 0.89 | 0.92 | 2 | 2 | Rigid |
| **3LVK** | 2.26 | 1.76 | 0.55 | 0.73 | 1 | 2 | Rigid |
| **3MJ9** | 8.22 | 5.11 | 0.02 | 0.13 | 0 | 0 | Rigid |
| **3MXW** | 12.45 | 11.22 | 0.00 | 0.00 | 0 | 0 | Rigid |
| **3P57** | 12.61 | 1.85 | 0.00 | 0.96 | 0 | 2 | Rigid |
| **3PC8** | 1.41 | 1.32 | 1.00 | 1.00 | 2 | 2 | Rigid |
| **3RVW** | 11.04 | 5.64 | 0.00 | 0.14 | 0 | 0 | Rigid |
| **3SE8** | 2.58 | 2.61 | 0.62 | 0.62 | 1 | 1 | Rigid |
| **3SGQ** | 1.56 | 1.54 | 0.83 | 0.83 | 2 | 2 | Rigid |

| PDB | Interface RMSD | | $f_{nat}$ | | CAPRI Rank | | Flexibility |
|---|---|---|---|---|---|---|---|
| | AF2 | AlphaRED | AF2 | AlphaRED | AF2 | AlphaRED | |
| **3U7Y** | 2.17 | 2.07 | 0.86 | 0.89 | 1 | 1 | Rigid |
| **3VLB** | 1.13 | 0.98 | 0.87 | 0.90 | 2 | 3 | Rigid |
| **3WD5** | 11.37 | 11.59 | 0.00 | 0.00 | 0 | 0 | Rigid |
| **4CPA** | 2.07 | 1.96 | 0.56 | 0.69 | 1 | 2 | Rigid |
| **4DN4** | 12.22 | 12.14 | 0.00 | 0.00 | 0 | 0 | Rigid |
| **4FP8** | 15.80 | 5.92 | 0.00 | 0.17 | 0 | 0 | Rigid |
| **4FQI** | 18.79 | 18.38 | 0.00 | 0.00 | 0 | 0 | Rigid |
| **4G6J** | 11.68 | 3.36 | 0.00 | 0.25 | 0 | 1 | Rigid |
| **4G6M** | 7.13 | 3.01 | 0.03 | 0.27 | 0 | 1 | Rigid |
| **4GXU** | 18.48 | 18.34 | 0.00 | 0.00 | 0 | 0 | Rigid |
| **4H03** | 2.13 | 2.01 | 0.43 | 0.64 | 1 | 1 | Rigid |
| **4HX3** | 15.67 | 15.51 | 0.00 | 0.00 | 0 | 0 | Rigid |
| **4M5Z** | 1.16 | 1.40 | 0.88 | 0.84 | 2 | 2 | Rigid |
| **4M76** | 19.94 | 4.24 | 0.00 | 0.15 | 0 | 0 | Rigid |
| **4Y7M** | 4.94 | 3.80 | 0.24 | 0.34 | 0 | 1 | Rigid |
| **5GRJ** | 13.59 | 3.60 | 0.04 | 0.35 | 0 | 1 | Rigid |
| **5JMO** | 15.48 | 5.05 | 0.00 | 0.14 | 0 | 0 | Rigid |
| **5O14** | 18.13 | 3.80 | 0.00 | 0.17 | 0 | 1 | Rigid |
| **5O1R** | 1.24 | 1.18 | 0.86 | 0.90 | 2 | 2 | Rigid |
| **5SV3** | 1.29 | 1.26 | 0.89 | 0.92 | 2 | 2 | Rigid |
| **5WK3** | 14.82 | 8.13 | 0.00 | 0.04 | 0 | 0 | Rigid |
| **5WUX** | 11.25 | 4.51 | 0.00 | 0.31 | 0 | 0 | Rigid |
| **5X0T** | 1.27 | 1.32 | 0.94 | 0.89 | 2 | 2 | Rigid |
| **5Y9J** | 7.09 | 2.76 | 0.07 | 0.42 | 0 | 1 | Rigid |
| **6A77** | 1.99 | 1.90 | 0.61 | 0.66 | 2 | 2 | Rigid |
| **6B0S** | 1.65 | 1.55 | 0.80 | 0.84 | 2 | 2 | Rigid |
| **6BPC** | 4.08 | 3.67 | 0.09 | 0.14 | 0 | 1 | Rigid |
| **6CWG** | 2.98 | 1.74 | 0.50 | 0.73 | 1 | 2 | Rigid |
| **6DBG** | 9.28 | 2.90 | 0.00 | 0.32 | 0 | 1 | Rigid |
| **6OC3** | 13.41 | 4.32 | 0.04 | 0.17 | 0 | 0 | Rigid |
| **7CEI** | 1.26 | 1.33 | 0.85 | 0.93 | 2 | 2 | Rigid |

#### References.
